# The intracellular domain of the epilepsy protein PCDH19 regulates spine density in cortical neurons *in vivo* via *Xlr* genes

**DOI:** 10.1101/2023.10.25.563961

**Authors:** Sylvia A Newbold, Victoria Becerra-Espinosa, Jaime Fabra-Beser, Ian WJ Fox, Cristina Llinares-Benadero, Elizaveta Stebleva, Cristina Gil-Sanz, Isabel Martinez-Garay

## Abstract

Mosaic mutations in the X-linked cell adhesion molecule Protocadherin 19 (*PCDH19*) lead to epilepsy with cognitive impairment, whereas complete absence of functional protein does not elicit any symptoms. It is believed that mosaic expression of PCDH19 leads to defective neuronal communication, but whether further roles beyond cell adhesion are critical for PCDH19 function in the cortex is currently unknown. We confirm that the proteolytic processing of PCDH19, previously described in hippocampal neurons, also takes place in cortical neurons *in vivo* and show that nuclear transport of its intracellular domain is mediated by importins. RNAseq analysis further indicates that the intracellular domain of PCDH19 leads to broad transcriptomic changes. Finally, we use *in utero* electroporation to provide the first *in vivo* data about the role of this cleaved intracellular domain in upper layer cortical neurons, where it reduces spine density through an increase in *Xlr* gene expression without affecting overall dendritic morphology. Our results suggest that PCDH19 could act as an activity sensor in a synapse to nucleus signalling pathway involved in synaptic homeostasis.

## INTRODUCTION

Cell adhesion molecules (CAMs) play essential roles in the development and functioning of the nervous system. Their ability to mediate homo- or heterophilic binding across the synaptic cleft allows them to connect pre- and postsynaptic neurons and underpins their important function in the formation of new synapses during development. For example, CAMs act as recognition molecules and coordinate synaptic differentiation, both morphologically and molecularly (Dean et al., 2003; Wit and Ghosh, 2016). In the adult brain, CAMs can regulate different aspects of synaptic structure and function, including spine shape, receptor function and synaptic plasticity (Dalva et al., 2007; Thalhammer and Cingolani, 2014). One way in which CAMs accomplish this is through direct interaction with neurotransmitter receptors and scaffolding proteins to modulate receptor surface levels (Nuriya and Huganir, 2006; Saglietti et al., 2007; Fièvre et al., 2016; Bassani et al., 2018).

Additionally, the strength of the synapses can be altered by processing through the physical loss of homo- or hetero-interactions across the synaptic cleft (Peixoto et al., 2012; Servián-Morilla et al., 2018). Finally, proteolytic processing of CAMs can translate neuronal activity into intracellular signalling via activity dependent proteolytic cleavage (Shinoe and Goda, 2015). This phenomenon, in which CAMs are processed by proteases in response to neuronal activity, can activate or suppress CAM-dependent signalling and generate new bioactive fragments with high spatial specificity restricted to active synapses. Numerous CAMs belonging to different families, including cadherins, have been shown to undergo activity dependent proteolytic cleavage (Nagappan-Chettiar et al., 2017). In the cadherin superfamily examples include N-cadherin (CDH2), PCDHγC3 and Protocadherin 19 (PCDH19) (Reiss et al., 2006; Marambaud et al., 2003; Gerosa et al., 2022). However, the relative contribution of adhesive and nuclear functions to the *in vivo* roles of these CAMs is still unknown.

PCDH19 is a member of the cadherin superfamily of cell-cell adhesion proteins (Wolverton and Lalande, 2001). Mutations in this X-linked gene lead to epileptic encephalopathy in heterozygous females and in males with somatic mutations, but do not cause any symptoms in hemizygous males (Dibbens et al., 2008; Juberg and Hellman, 1971; Depienne et al., 2009). This unusual inheritance pattern is believed to be due to a phenomenon called “cellular interference”, in which the coexistence of cells with different genotypes, caused by random X chromosome inactivation in heterozygous females and by somatic mutation in males, is detrimental at the tissue level, even if mutated cells are not affected themselves (Depienne et al., 2009). PCDH19 can mediate cell adhesion on its own, but also as a complex with CDH2 (Emond et al., 2011) or other delta protocadherins (Bisogni et al., 2018; Pederick et al., 2018). In addition, PCDH19 has been shown to localize to synapses and to interact with the alpha subunit of the GABA receptor (Bassani et al., 2018; Hayashi et al., 2017), modulating surface availability and affecting postsynaptic inhibitory currents. PCDH19 mosaicism has been shown to influence mossy fiber-CA3 synaptic function, plasticity and behaviour in mice (Hoshina et al., 2021).

Beyond its roles at the membrane, PCDH19 has also been reported to interact with the paraspeckle protein NONO and co-regulate oestrogen receptor alpha controlled genes (Pham et al., 2017), and to regulate expression of immediate early genes after proteolytic cleavage (Gerosa et al., 2022).

However, the transcriptional pathways regulated by PCDH19 and the *in vivo* consequences of this nuclear function remain unclear. Thus, to better understand the biological significance of PCDH19 non-adhesive functions in cortical neurons, we verified its processing in these cells and carried out a transcriptional analysis of the changes triggered by its intracellular domain (ICD). Importantly, we conducted *in vivo* functional studies in the mouse neocortex, revealing that PCDH19-ICD regulates spine density in upper layer cortical neurons through the upregulation of *Xlr* gene expression.

## MATERIALS AND METHODS

### Animals

C57BL/6J WT mice were purchased from Charles River Laboratories. *Pcdh19* knock-out (KO) mice (TF2108) were purchased from Taconic Biosciences. Ai9 Cre reporter strains (Madisen et al., 2010) were originally obtained from the Jackson laboratory and are currently bred in the animal facilities of the University of Valencia, placed in the Burjassot campus. Animals were housed on a 12 h light/dark cycle and with *ad libitum* access to food and drink. All procedures were conducted in accordance with the Animals (Scientific Procedures) Act1986 (amended 2012). Genotyping of the *Pcdh19* KO animals was done using the Mouse Direct PCR kit (Biotool, cat no. B4001), following the manufacturer’s instructions and using primers *Pcdh19*-WT-F (5’-TAGAGGTTCTTGCTGAAGACTTCC-3’), *Pcdh19*-WT-R (5’-TCAACTGTTTCGATGAGACACTGC-3’), *Pcdh19*-Mut-F (5’-GTGCGTACCAGGCGGGAGC-3’) and *Pcdh19*-Mut-R (5’-CCCTAGGAATGCTCGTCAAGA-3’). Ai9 animals were genotyped following protocol 29436 from the Jackson Laboratory.

### Plasmids

pMet7-PCDH19-Gal4DBD-VP16 was created by In-Fusion cloning (Takara Bio) using PCR products for PCDH19-FL, Gal4DBD-VP16 and a pMet7 backbone. Gal4DBD-VP16 fragment was amplified from Gal4-VP16, a gift from Lea Sistonen (Addgene plasmid # 71728) (Budzyński et al., 2015). pGL2-GAL4-UAS-Luc was a gift from Martin Walsh (Addgene plasmid # 33020) (Nishio and Walsh, 2004) pMet7-IFNAR1-Gal4-VP16 and pMet7-IFNAR2-Gal4-VP16 plasmids (the controls for the cleavage luciferase reporter assay) were kind gifts from the Tavernier lab, Ghent, Belgium and pRL-TK was acquired from Promega.

pCAGIG (pCIG) was a gift from Connie Cepko (Addgene plasmid # 11159) (Matsuda and Cepko, 2004) and pCAG-PCDH19FL-HA contains full length mouse PCDH19 (including exon 2) fused C-terminally with an HA tag under the control of the Chicken β-actin promotor. pCAG-PCDH19ICD-HA is the same plasmid but contains PCDH19 aa 700 to 1145 with an added methionine at the beginning.

To create pCAG-PCDH19ICD(NLSmut)-HA, the last four basic residues of the predicted NLS (aa 777 to 780 of the FL protein) were mutated via PCR from “KKKK” to “AAAA” in the PCDH19-ICD-HA construct. KPNA1-myc was purchased from Origene (CAT#: MR208599). pCAG-PCDH19ICD-HA-i-Cre was created by In-Fusion cloning (Takara Bio) using PCR products for IRES, Cre and the pCAG-PCDH19ICD-HA backbone. pCBA was derived from pCAGIG by eliminating the IRES-EGFP sequences through digestion and re-ligation.

To create pZDRosa-floxedNeo-Pcdh19-CYTO-HA (the targeting vector for PCDH19-ICD-HA overexpression from the *Rosa26* locus), the pZDRosa-floxedNeo-IRES-EGFP plasmid (kind gift of Dr Xinsheng Nan) was linearized and the IRES-EGFP fragment was excised by double restriction digestion with BsrGI-HF and AscI. The PCDH19ICD-HA fragment was amplified from the pre-existing plasmid pCIG-19ICD-HA using primers Rosa26-CYTO-HA-F2 (5’-ACCTCGAGTGGCGCGCCGCGCAGCCATGGCAATGGCAATCAAATGC −3’) and Rosa26-CYTO-HA-R (5’-CCGCTTTACTTGTACTCAAGCGTAATCTGGAACATCGTATG −3’).

The resulting PCR product was cloned between the 3’ and 5’ arms of the *Rosa26* targeting vector using In-Fusion cloning (Takara Bio). Plasmid sequence was checked by sequencing. pCMV-RosaR4 KKR mutations and pCMV-RosaL6 ELD mutations were gifts from Charles Gersbach (Addgene plasmids # 37199 and # 37198) (Perez-Pinera et al., 2012).

shRNA plasmids against *Xlr* genes were a kind gift of Dr Marta Nieto (Cubelos et al., 2010).

### Cell culture, transfection and drug treatment

Cells in culture were routinely tested for mycoplasma infection with Lookout Mycoplasma PCR detection kit (Sigma, MP0035), following the manufacturer’s instructions. HEK293 and HeLa cells were maintained in CA media (DMEM + 1% non-essential amino acids + 1% L-Glutamine + 10% FBS heat inactivated + 1.43 mM-β-mercaptoethanol) on 100 mm (Nunc) dishes. For 6 well plates, ∼500,000 cells were split into each well. For 12 well and 24 well plates, ∼250,000 and ∼50,000 cells were split into each well, respectively. Cells were transfected 24 hours after seeding using Lipofectamine™ 2000 (Thermo Fisher) at a 1:2 ratio (DNA: Lipofectamine). 1 μg and 500 ng of DNA were used for each plasmid for co-immunoprecipitation (CoIP) and immunocytochemistry (ICC) experiments, respectively.

Cells (HeLa cells and/or mESC-derived neurons) were subjected to the following treatments: Ionomycin (Sigma, I3909) 5 μM for 10 min, 30 min or 60 min; GI254023X (Sigma, SML0789) 10 μM for 1h; MK-8931 (Stratech, B6195-APE) 10 μM for 1h; NMDA (Sigma, M3262) 50 μM for 30 min and (+)-MK-801 maleate (Tocris, 0924) 1 μM for 30 min.

### Cleavage luciferase reporter assay

HEK293 cells (5×10^4^ HEK293T cells/well in a 24-well plate) were transfected with 150 ng pGL2-Gal4-UAS-Luc, 5 ng pRL-TK and 150 ng of the appropriate Gal4-VP16 fused constructs: pMet7-PCDH19-Gal4-VP16, pMet7-IFNAR1-Gal4-VP16 and pMet7-IFNAR2-Gal4-VP16 using Lipofectamine™ 3000. Firefly and Renilla luminescence activities were measured one day after transfection using the Dual Luciferase Reporter Assay System (Promega), following the passive cell lysis protocol.

Measurements were performed with a Victor3 1420 Multilabel Counter (Perkin Elmer) and the ratio of Firefly to Renilla values was expressed as relative fluorescence units. Experiments were performed in two technical and six biological replicates.

### Cell lysis for RNA/protein extraction

For RNA extractions, neurons were rapidly washed twice with 1X PBS and lysed in 350 μl of cold RLT buffer (Qiagen) with 1% β-mercaptoethanol and collected immediately on ice. RNA sequencing samples were rapidly transferred and stored at −80°C until all samples were collected. For protein extractions, cells were rapidly washed twice with 1X PBS and lysed in 100 μl of RIPA buffer freshly supplemented with protease and phosphatase inhibitors (50 mM Tris-HCl, 150 mM sodium chloride, 1 mM EDTA, 1% triton X-100, 0.2% sodium deoxycholate supplemented with: 1.5 mM aprotinin, 100 mM 1-10 phenantroline, 100 mM 6-aminohexanoic acid, 1% protease inhibitor cocktail (Sigma), 1% phosphatase inhibitor cocktail (Sigma)). Lysates were kept on ice for 30 minutes with brief vortexing every 5 minutes. Samples were then centrifuged at 14000g for 10 minutes and supernatant was transferred to a clean tube. Samples were aliquoted and stored at −80°C until used for western blotting.

### Immunoprecipitation

For immunoprecipitation, cells or tissue were lysed in freshly made IP lysis buffer (20 mM Tris-HCl, 150 mM NaCl, 1mM EDTA, 1% Triton X, 10 mM NaF, 1mM Na3VO4,1% protease inhibitor cocktail (Sigma), 1% phosphatase inhibitor cocktail (Sigma)). 10 μl of Protein G Sepharose beads were washed twice with 500 μl of cold 1X PBS by centrifugation (2000g, 2 min, 4°C). In parallel, tissue or cell-lysate samples were centrifuged (14000g, 10 min, 4°C). Sample supernatant was pre-cleared with the washed beads for 30 min at 4°C under constant rotation. Beads and non-specifically bound proteins were precipitated by centrifugation (2000g, 2 min, 4°C). 10% of sample supernatant was put aside and saved to be used as INPUT control. The remaining 90% of the supernatant was used for immunoprecipitation and added to 20 μl of pre-washed Protein G Sepharose beads with 2 μl of antibody of interest. After a 2-hour incubation at 4°C with constant rotation, samples were centrifuged (2000g, 2 min, 4°C) and washed with lysis buffer (3X 2000g, 2min, 4°C). Finally, samples were eluted in LDS buffer and incubated for 10 min at 70°C). Finally, the beads were removed by centrifugation (2000g, 5 min, RT). Samples were stored at −80°C until analysed by western blot. The antibodies used for co-IP were anti-MYC (MA1-980 mouse monoclonal, Thermo Fisher), anti-KPNA1 (18137-1-AP rabbit polyclonal, Proteintech) and anti-PCDH19 (A304-648A rabbit polyclonal, Bethyl).

### Western blotting

Protein lysates were prepared by addition of LDS buffer and 10% 0.5 M DTT, and boiled at 70°C for 10 min. Samples were then centrifuged at 14000 g for 10 min and loaded onto a NuPAGE Novex 4-12% Bis-Tris gel (Novex Life Technologies, WC1020) and run at 120 V for 90 minutes.

Proteins were transferred to a nitrocellulose membrane with a 0.2 μm pore size (GE Healthcare Life Sciences, 10600001) by wet transfer at 100 V for 120 minutes. Membranes were incubated shaking for 1 hour at RT with 4% blocking solution (5% milk powder (BioRad) in TBS-T). Primary antibody incubation was done overnight at 4°C shaking. The following day, membranes were washed 3 times for 10 minutes in TBS-T and then incubated for 1 hour at RT with the appropriate secondary antibody (in 5% milk powder in TBS-T blocking). Membranes were washed again 3 times for 10 minutes in TBS-T. Blots were finally developed with 1 ml of WesternBright ECL substrate (Advansta) and imaged with a ChemiDoc XRS+ (BioRad), using the Image Lab software.

For western blot analysis of tissue samples, the protein concentration of each sample was measured using the Micro BCA™ Protein Assay Kit (Thermo Fisher), following the manufacturer’s instructions. Colorimetric intensity was measured using a FLOUstar Omega microplate reader (BMG Labtech). The samples were then diluted in lysis buffer to reach a concentration of 40 μg which was then used for western blotting as described above.

Primary antibodies used for western blot include: anti-PCDH19 C-terminal (1:1000, A304-648A rabbit polyclonal, Bethyl); anti-N-Cadherin (1:1000, 33-3900 mouse monoclonal, clone 3B9, Thermo Fisher); anti-Pan-Cadherin (1:1000, ab6529 rabbit polyclonal, Abcam); anti-HA (1:2000, ROHAHA rat polyclonal, clone 3F10, Roche); anti-MYC (1:2000, MA1-980 mouse monoclonal, Thermo Fisher); anti-Histone H3 (1:5000, ab1792, rabbit polyclonal, Abcam) and anti-β-Actin (1:2000, ab8226 mouse monoclonal, Abcam). Secondary antibodies used for western blot include anti-Rabbit-HRP (1:20000, Promega W4011); anti-Mouse-HRP (1:20000, Promega W4021) and anti-Rat-HRP (1:20000, R&D systems HAF005).

### Subcellular fractionation

Mouse embryonic fibroblast (MEF) cell lysates were processed using the Mem-PER™ Plus Membrane Protein Extraction Kit (Thermo Fisher, 89842) following the manufacturer’s instructions.

### IHC and ICC

Animals were injected with 100 μl of Euthatal (Merial, R02701A) and transcardially perfused with 30 ml of 1X PBS, followed by 30 ml of 4% PFA. Brains were postfixed in 4% PFA overnight at 4°C, then washed in 1X PBS the following day and stored at 4°C in the dark until sectioned. 150 μm P30 brain sections, or cells on glass coverslips, were washed in 1X PBS for a minimum of 3 times, followed by several washes in PBS 1X containing 0.25% triton X-100 (0.25% PBS-T). Sections, or cells were incubated at RT for at least 3 hours in BSA/blocking solution in 0.25% PBS-T, then incubated with the primary antibodies overnight at 4°C in the dark. The following day, sections or cells were washed, and incubated with appropriate fluorescently conjugated secondary antibodies, washed, counterstained with DAPI (1:4000 in 1X PBS) and mounted with DAKO mounting media or ProLong™ Diamond Antifade Mountant (Invitrogen) on glass slides. For immunostaining of P60 electroporated brains, 150 μm sections were blocked in PBS-T 0.5% and 10% Horse Serum for 5 hours, followed by overnight (16h) incubation with primary antibodies at room temperature under constant shaking. After that time, sections were thoroughly washed (3 x 30 min) with 0.5% PBS-T and incubated with the appropriate fluorescently conjugated secondary antibodies for 5 hours at RT, washed, counterstained with DAPI (1:4000 in 1X PBS) and mounted with DAKO mounting media on glass slides.

Primary antibodies used for immunostaining included anti-HA (1:500, ROHAHA rat polyclonal, clone 3F10, Roche) and anti-GFP (1:2000, A10262 chicken polyclonal antibody, Thermo Fisher; 1:1000, 600-101-215M goat polyclonal antibody, Rockland). Secondary antibodies included: anti-chicken 488 (Thermo Fisher; A11039 Alexa Fluor® 488 goat anti-chicken; 1:1000) and anti-goat 488 (Jackson ImmunoResearch, Alexa Fluor® 488 donkey anti-goat #705-545-003; 1:1000).

### Neuronal differentiation of mESCs

E14 male mouse embryonic stem cells (mESCs) used in this study were kindly provided by Dr. Xinsheng Nan (Cardiff University). Differentiation into cortical-like neurons was done following the protocol by (Bibel et al., 2004) (Bibel et al., 2007). Although E14 mESCs are feeder-independent, in some instances, feeder passaging was added to the differentiation protocol to improve their quality. 12-well plates were used for protein and RNA extraction and 4-well plates for transfections and immunocytochemistry. Cells were plated in a range between ∼750,000 and ∼1.5*10^6^ cells/well, depending on downstream applications.

### Genetic engineering of mESCs

E14 mESCs were passaged the day before nucleofection. On the day, they were trypsinized and 4*10^6^ cells were used for one round of nucleofection. In brief, cells were pelleted and resuspended in 100 μl of P3 transfection solution (82 μl Amaxa Buffer and 18 μl P3 supplement; Lonza) and 10 μl of DNA mix. The following amount of plasmids was used: 10 μg of the targeting construct (pZDRosa-floxedNeo-Pcdh19-CYTO-HA) and 1 μg each of the two zinc finger nuclease (ZFN) plasmids: pCMV-RosaR4 KKR mutations, containing the right ZFN (ZFN-R), and the pCMV-RosaL6 ELD mutations, containing the left ZFN-L (ZFN-L). Cells were nucleofected using the 4D-Amaxa Nucleofector X-unit (Lonza) and the CG104 programme. Immediately after nucleofection cells were plated at low density for antibiotic selection. For removal of the neomycin resistance cassette, 10 μg of the pCIG-CRE plasmid was nucleofected as described above. After ZFN targeting, nucleofected cells were suspended in 10 ml of ESC medium and plated at densities ranging between 0.625 and 2.5 ml/10 cm dish. Cells underwent a 10-day selection process with 250 μg/ml of G418 (Geneticin), with media changed every two days. After about 10 days, or when they were visible by the naked-eye, 100 colonies were manually picked. In brief, cells were incubated for a couple of minutes with 0.01% trypsin (0.05% trypsin, diluted in PBS) in order for colonies to detach from the plate but without dissociating. Colonies were then carefully transferred to individual wells in a 96-well plate, trypsinized with 0.05% trypsin, resuspended and transferred to a 24-well plate to grow. Clones were expanded for DNA and protein extraction and then frozen. For mESC subcloning, after nucleofection for removal of the selection cassette, cells were plated at a density of 300 cells/10 cm plate. 24 colonies originating from different clones were picked as described above. Once expanded, these clones were “reverse selected” to test out loss of antibiotic resistance.

### mESC genotyping

Cells were pelleted, resuspended in 500 μl of cell lysis buffer (10 mM Tris, pH 8.0, 1 mM CaCl2; 100 mM NaCl; 0.5% SDS; 5 mg/mL proteinase K (Promega)) and incubated overnight at 50°C. The following day, 500 μl of 100% isopropanol and 50 μl of 3 M NaOAc were added to precipitate DNA. DNA was pelleted by centrifugation (15 minutes at top-speed), washed with 70% ethanol and resuspended in 30 μl of TE buffer (Qiagen). Clones were genotyped by PCR, using the long-range S equalPrep PCR kit (Thermo Fisher) and primers ReverseR26OUT2 (5’arm genomic; 5’-CAAGCGGGTGGTGGGCAGGAATGCG-3’), Neo-pR2 (5’ arm selection cassette; 5’-TCGGCAGGAGCAAGGTGAGATGAC-3’), ForwardR26OUT2 (3’ arm genomic; 5’-ACCAGAAGAGGGCATCAGATCCCATTAC-3’) and gen19-ICDF2 (3’ arm intracellular domain; 5’-GCGTGAAGCGTCTGAAGGATATCGTTC-3’).

### Karyotyping

mESC clones to be karyotyped were incubated with demecolcine solution (0.1 μg/ml) for two hours in the incubator. Cells were then trypsinized, pelleted and washed by centrifugation in 1X PBS twice, and subsequently resuspended in 2 ml of 1X PBS and 6 ml of hypotonic 0.0375 M potassium chloride solution and incubated for 12 min at 37°C. Cells were then pelleted, supernatant removed and a 3:1 volume:volume ratio of cold methanol/acetic acid mixture (−20°C) was added dropwise.

After a 20 min incubation at room temperature, cells were pelleted again, supernatant removed, and fresh methanol/acetic acid was added. Cells were centrifuged one last time, the supernatant was removed, this time leaving about 100 μl, in which the cells were resuspended. Finally, cells were dropped from about 20 cm height on glass slides. Slides were left to dry, stained with DAPI (1:4000 in ddH2O), coverslipped and imaged immediately. A minimum of 10 cells were imaged for each clone. Chromosomes were then counted using the ImageJ (Fiji) cell-counter plug-in.

### RNA sequencing

RNA extraction was done with RNeasy Kit (Qiagen) in RNase-free conditions following the user’s manual with DNase treatment (Qiagen). Quality control of the samples was done via Tapestation (Agilent Technologies) and RNA integrity number (RIN) was determined for all samples. Concentration of samples was measured by QUBIT. RNA sequencing was done at Cardiff University Genomic Hub. Libraries were prepared following Illumina’s TruSeq Stranded mRNA sample preparation guide. In brief, mRNA was purified from total RNA using poly-T oligos, mRNA was then fragmented into smaller fragments and random priming was used for cDNA synthesis. The sequencing was carried out on an Illumina Nextseq 500 platform with 4 cartridges PE (2×75bp) sequencing on high output 150 cycle V2.5 cartridges. 1% Phix was spiked into each run as per the Illumina recommendations. The samples were pooled to obtain equal reads for each sample with an aim of at least 44 M reads per sample. Sequencing was paired end. Quality control of sequencing run, such as QC content and sequence duplication was performed before downstream analysis. Paired end reads from Illumina sequencing were trimmed of adaptor sequences with Trim Galore and assessed for quality using FastQC, using default parameters. Reads were mapped to the mouse GRCm38 reference genome using STAR (Dobin et al., 2013) and counts were assigned to transcripts using featureCounts (Liao et al., 2014) with the GRCm38 Ensembl gene build GTF. Both the reference genome and GTF were downloaded from the Ensembl FTP site (http://www.ensembl.org/info/data/ftp/index.html/). Differential gene expression analyses used the DESeq2 package (Love et al., 2014), using the Benjamini-Hochberg correction for multiple testing. Differential gene-splicing analyses used the DEXSeq package (Anders et al., 2012), (Reyes et al., 2013) also using the Benjamini-Hochberg correction for multiple testing.

### R packages for plotting

All plotting of RNA sequencing data was done on R (v.4.02) via RStudio (v.1.2.1335).

Plotting was done using R package “ggplot2” (v.3.3.2). Over-representation analysis and Gene Set Enrichment analysis was done via “clusterProfiler” (v.3.16.1) (Yu et al., 2012).

### Quantitative real-time PCR

For RNA extraction, samples were collected and protected with RNAlater (Thermo Fisher) at −80° C for RNA extraction, which was performed using RNeasy Mini Kit (Qiagen) followed by RNase-Free DNase set (Qiagen). Maxima First Strand cDNA Synthesis Kit was used to generate the cDNA template for quantitative real-time PCR (Thermo Fisher). RT-PCRs were carried out with Applied Biosystems StepOne Plus and analysed using the corresponding software StepOne Software Version 2.0 (Applied Biosystems). Quantification was performed using a standard curve. The primers used included *Erbb4*-fw (5’-CAAAGCCAACGTGGAGTTCATGG-3’), *Erbb4*-rv (5’-CTGCGTAACCAACTGGATAGTGG-3’), *Lhx2*-fw (5’-GATGCCAAGGACTTGAAGCAGC-3’), *Lhx2*-rv (5’-TTCCTGCCGTAAAAGGTTGCGC-3’), *Zic1*-m-fw (5’-TTTCCTGGCTGCGGCAAGGTTT-3’) and *Zic1*-m-rv (5’-ACGTGCATGTGCTTCTTGCGGT-3’).

### *In utero* electroporation

For electroporations plug checking was performed, with noon of the day the plug was found considered as embryonic day 0.5 (E0.5). Timed-pregnant females were deeply anesthetized with isoflurane and maintained in 1.5-2% isoflurane during the surgery. The abdominal cavity was opened, and the uterine horns were exposed. The DNA solution was injected into the lateral ventricle of the embryos through the uterus wall using pulled capillaries (PC-10, Digitimer). Embryos were electroporated at E15.5 by applying electric pulses (45 V; 85ms on/950 ms off/5 pulses) with an electric stimulator (BTX Electroporator ECM 830 (Harvard Apparatus)) using round electrodes (CUY650P5, NepaGene). DNA was diluted in 1X TE and coloured with 0.5% Fast Green (Sigma Aldrich). For PCDH19-ICD overexpression, the plasmids used were pCIG and pCIG-19ICD-HA at a concentration of 1 μg/ul. When electroporating Ai9 Cre reporter animals, we used pCIG-Cre and pCAG-PCDH19ICD-HA-i-Cre at 1 μg/ul. For rescue experiments, we used pCBA or pCIG-19ICD-HA at 1 μg/ul, combined with pCIG at 0.5 μg/ul and the corresponding shRNA plasmid at 1.5 μg/ul.

Embryos in each litter were electroporated with either control or 19ICD plasmids, using different hemispheres to differentiate them. All surviving pups were subsequently used for spine and axonal analyses, except when electroporation efficiency was too low to allow imaging. For RNAScope analysis we selected one or two electroporated animals per litter for each plasmid, based on electroporation efficiency. The results presented here are derived from a total of 37 animals.

Electroporations were performed by three and data analysis by four different researchers, so that for each dataset, at least one of the researchers carrying out data analysis was not involved in the electroporation experiments and did the analysis completely blind.

### RNAScope with IHC

Two RNAScope probes were designed. One recognizes *Xlr3a*, *Xlr3b*, *Xlr3c* and other *Xlr3*-like pseudogenes (such as *Xlr3d-ps* and *Xlr3e-ps*), and the other recognizes *Xlr4a*, *Xlr4b*, *Xlr4c* and other *Xlr4*-like pseudogenes, including *Xlr4d-ps* and *Xlr4e-ps*, due to their high sequence similarity. RNAScope was performed on 15 μm cryostat brain sections of P60 animals that had been electroporated with either pCIG-Cre or pCBA-PCDH19ICD-HA-i-Cre at E15.5 following the manufacturer’s instructions. Briefly, sections were washed for 5 min to remove the OCT and then dried for 5 min at 37°C. followed by 5 min at RT. Next, sections were postfixed in 4% PFA at 4°C for 15 min and dehydrated in 50%, 70% and 100% ethanol for 5 min each. After air-drying the slides for 5 min at RT, sections were treated with hydrogen peroxide 10 min at RT, rinsed and incubated in 100% EtOH again for another 5 min. After a 30 min treatment with Protease III at 40 °C, slides were rinsed twice in H2O and incubated with probes dilutes 1:50 in Probe Diluent Solution for 2 h at 40 °C. Slides were then washed twice with wash buffer at RT and stored o/n in 5X SSC at RT. The following day slides were washed twice and incubated with amplification solutions 1, 2 and 3 for 30 min each at 40°C. Next, slides were incubated with HRP solution (15 min at 40°C), followed by a 30 min incubation with 1:1000 640 TSA Vivid Fluorophore and a 15 min incubation with HRP Blocker at 40°C. Once the RNAScope was completed, slides were washed in PBS and incubated with blocking solution (10% horse serum, 0.2% Triton-X-100) for at least one hour at RT. Primary antibody incubation (in blocking solution) was performed o/n at 4°C and the following day slides were washed with PBS, incubated with secondary antibody diluted in blocking, stained with DAPI and mounted with ProLong Diamond (Thermo Fisher). Primary antibodies used for immunostaining included anti-HA (1:50, ROHAHA rat polyclonal, clone 3F10, Roche) and anti-GFP (1:1000, 1:1000, 600-101-215M goat polyclonal antibody, Rockland). Secondary antibodies included: anti-rat 488 (Thermo Fisher; A-21208 Alexa Fluor® 488 donkey anti-rat; 1:500) and anti-goat 488 (Jackson ImmunoResearch, Alexa Fluor® 488 donkey anti-goat #705-545-003; 1:1000).

### Microscopy

Brain sections or cells were imaged on a confocal microscope (Carl Zeiss, LSM 780, 800 or 980) with the Zen Black software (version 2.0, Carl Zeiss). For reconstruction of the neuronal morphology, 1 μm spaced Z-stack tiles, spanning the whole neuron, were taken with a 40X water-immersion objective. For spine analysis, representative segments of secondary order apical or basal dendrites belonging to the previously imaged neurons, were imaged with a 63X oil-immersion objective, as 0.5 μm spaced Z-stacks. Images of cultured cells (293HEKs, HeLa cells and mESC-derived neurons) were also taken with a 63X oil-immersion objective, as 1 μm spaced Z-stacks. For RNAScope stained sections, 1 μm spaced images were acquired using an Olympus FV10i confocal microscope. Low magnification (10X objective) and high magnification (60X oil-immersion objective) images of electroporated neocortices were taken for validation of RNAScope probe staining and for RNAScope dot quantification, respectively.

### Image Analysis

Densitometric analysis of western blots was done using Image Lab (v.6.0.1) (BioRad). Lanes and bands were drawn in the software and adjusted volume intensity of each band was extracted. When calculating proteolytic fragments, full-length protein and fragment band intensity were detected on the same blot but using different exposures to avoid saturation. Intensity of the proteolytic fragment was always calculated as a ratio of the full-length protein.

Neuronal tracing analysis was done using the Fiji plugin SNT (Arshadi et al., 2021). Spines were counted manually, and dendrite was traced with SNT to determine length and calculate spines/20 μm. Axonal analysis was performed using the Plot Profile tool in Fiji on a rectangular selection encompassing the axonal arborizations in the contralateral hemisphere of the electroporated brains. For RNAScope analysis, a single confocal plane was chosen and cells positive for anti-GFP or anti-HA were identified and their nuclei marked as regions of interest (ROIs). Those ROIs were then transposed to the far-red channel to count the number of dots from the RNAScope probe. For the rescue experiments, only neurons with somal HA signal were used for analysis.

### Experimental Design and Statistical Analysis

Statistical analysis was carried out on GraphPad Prism (version 10.3.1). Shapiro-Wilk test was used to test for normality of the data. Data sets that passed the normality test (P > 0.05) were analysed with two-tailed unpaired Student’s t-test when comparing two groups, or one-way ANOVA, followed by Tukey’s correction for multiple comparisons if more than 2 groups were compared. For data sets that failed the normality test, the non-parametric Mann Whitney test was performed instead to compare between two samples (none of the data sets used for multiple comparisons failed the normality test). Kernel density analysis was performed using the R density base function with a standard gaussian fit and a bandwidth of 1. For the migration analysis, a histogram distribution of the distance of each electroporated neuron to the top of the cortical plate was created first, using 50 μm bins and combining all neurons from each condition (pCIG: 7 brains, 1895 neurons; pCIG-19ICD-HA: 11 brains, 3045 neurons). Next, a Kolmogorov-Smirnov test was run between the two histograms. For spine density analysis, segments of 1-5 basal and apical dendrites were analysed per neuron and values were averaged, so that n represents individual neurons and not individual dendrites. Data are displayed as mean ± SEM in figures 1-6. However, in figure 1 data have been normalized to the values of one of the conditions for ease of visualization, even if the statistical analyses were performed with non-normalized data as described above.

**Figure 1.**
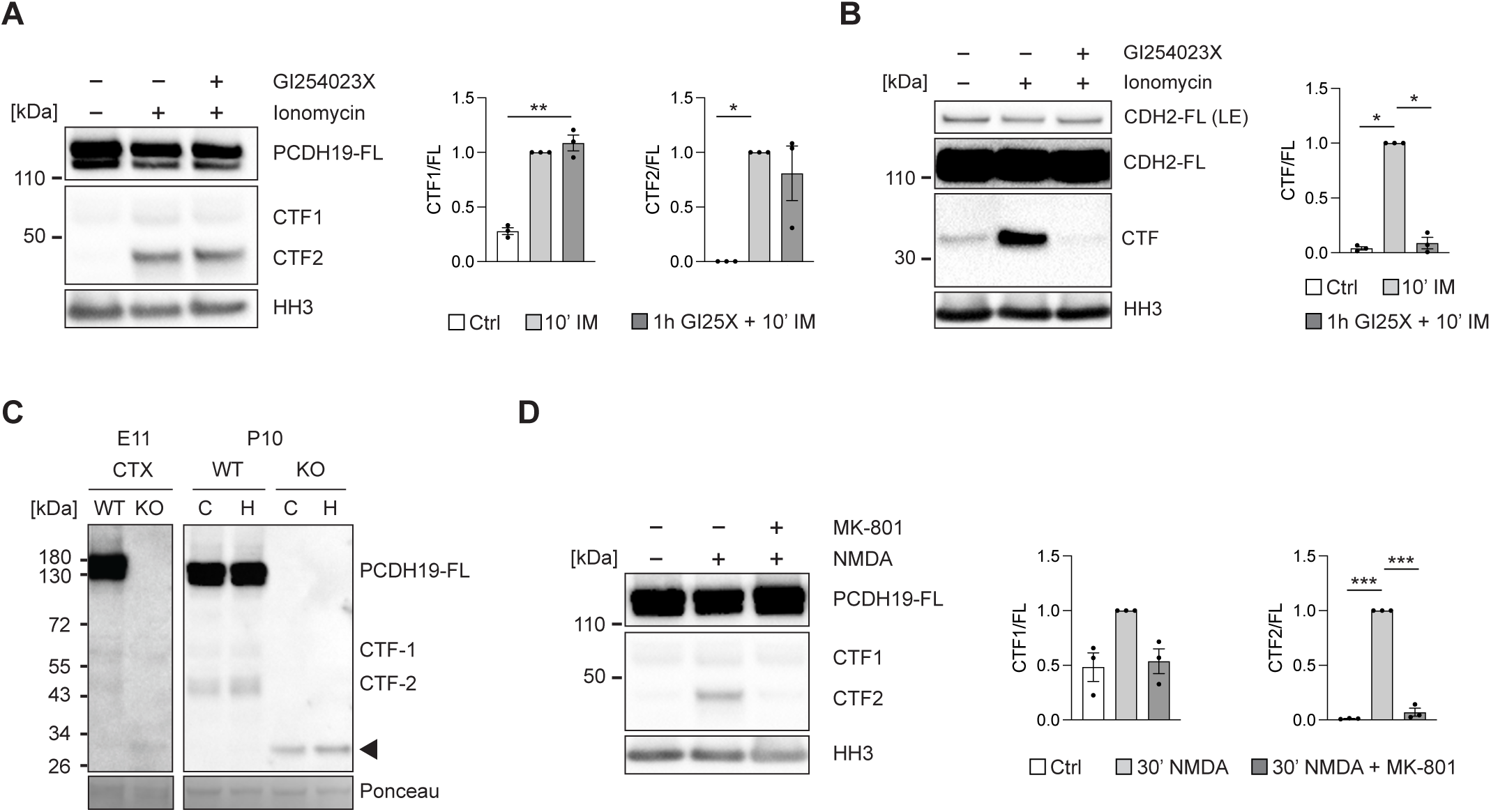
Proteolytic processing of PCDH19 in cortical neurons is not mediated by ADAM10 and is activity dependent. A. Western blot detection (left) and quantification (right) of full length and processed PCDH19 in mESC-derived neurons under control, ionomycin or ionomycin + GI254023X treatments. (N=3) B. Western blot detection (left) and quantification (right) of full length and processed CDH2 in mESC-derived neurons under control, ionomycin or ionomycin + GI254023X treatments. Same lysates as in panel A. (N=3) C. WB detection of PCDH19 in E11 cortical lysates and P10 cortical (CTX, C) and hippocampal (H) lysates of *Pcdh19* WT and KO mice. Arrowhead points to a ∼30kDa band present only in KO samples. D. Western blot detection (left) and quantification (right) of full length and processed PCDH19 in mESC-derived neurons under control, NMDA and NMDA + MK-801 treatments. (N=3) (A, B, D) The ratio of CTF to FL band intensity was used for quantification. Data are normalized to IM treatment (A, B) or NMDA treatment (D) for ease of visualization and are presented as mean ± SEM. N: Biological replicate. Relevant p-values are indicated: *p < 0.05, **p < 0.01, ***p < 0.001. Statistical analysis: one-way ANOVA with Tukey’s post-hoc test on non-normalized values.

Axonal analysis was carried out by fitting a mixed model to the normalized data, considering two variables: distance (1-100) and plasmid (pCIG-Cre vs pCIG-19ICDHA-i-Cre). Before that, and to account for small variations in cortical thickness between brains, average fluorescence intensity had been calculated for each of 100 bins spanning from the pial surface to the white matter.

## RESULTS

### Processing of Protocadherin 19 in cortical neurons is independent of ADAM10

Processing of PCDH19 in cultured hippocampal neurons has been previously described (Gerosa et al., 2022). We independently confirmed proteolytic processing of this adhesion protein in HEK293 cells using a luciferase reporter assay (Fig. S1A-C) and investigated the processing of PCDH19 in cortical neurons by treating ESC-derived cortical-like neurons, obtained following an established protocol that yields a highly pure (95%) population of these cells (Bibel et al., 2007), with ionomycin (a calcium ionophore previously described to increase proteolytic processing of cadherin family members (Reiss et al., 2005; Marambaud et al., 2002)). This treatment greatly increased the intensity of two smaller bands above and below 50 kDa, which match the size of the predicted membrane bound cytoplasmic fragment 1 (CTF1) and soluble CTF2 fragments (Fig. 1A). These results demonstrated that PCDH19 undergoes proteolytic processing in cortical-like neurons and that the main product of that cleavage is the CTF2 fragment of about 45 kDa.

ADAM10 functions as sheddase for several members of the cadherin superfamily (Reiss et al., 2005; Maretzky et al., 2005; Reiss et al., 2006; Bouillot et al., 2011) and has also been reported to be involved in PCDH19 processing in hippocampal neurons (Gerosa et al., 2022). Thus, we hypothesized that PCDH19 might also be cleaved by this protease in our ESC-derived neurons and we set out to explore this possibility using the ADAM10 specific inhibitor GI254023X. However, a one-hour treatment with GI254023X before ionomycin addition did not reduce the intensity of the CTF1 and CTF2 bands (CTF1, Ctrl: 0.3 ± 0.07, 10’ IM: 1.32 ± 0.13, 1h GI25X + 10’ IM: 1.35 ± 0.22; n = 3; one way ANOVA, F(2,6) = 14.84, *p* = 0.0048; CTF2, Ctrl: 0.0005 ± 0, 10’ IM: 0.24 ± 0.03, 1h GI25X + 10’ IM: 0.2 ± 0.08; n = 3; one way ANOVA, F(2,6) = 6.905, *p* = 0.0278; Fig. 1A). To confirm that our treatment was effectively inhibiting ADAM10, we blotted the same lysates with an antibody against CDH2, which revealed a significant decrease in the intensity of its 37 kDa CTF band, confirming ADAM10 inhibition and a lack of involvement of ADAM10 in PCDH19 processing in cortical-like ESC-derived neurons (Ctrl: 0.07 ± 0.02, 10’ IM: 1.97 ± 0.52, 1h GI25X + 10’ IM: 0.23 ± 0.16; n = 3; one way ANOVA, F(2,6) = 11.30, *p* = 0.0092; Fig. 1B). Our results indicate that proteolytic processing of PCDH19 also takes place in cortical-like neurons but that, unlike in cultured hippocampal cells, matrix metalloproteases or A Disintegrin and metalloproteinase domain-containing proteins other than ADAM10 act as the main sheddases here. These data suggest that the proteases involved in PCDH19 processing vary between different neuronal types.

### Protocadherin 19 is processed in the cortex *in vivo*

We sought to confirm that proteolytic processing of PCDH19 also happens in the cortex *in vivo*. Western blot analysis of PCDH19 expression in mouse embryonic forebrain (E11) and postnatal day 10 (P10) cortical and hippocampal lysates detected CTF1 and CTF2 bands in wild type (WT) tissue, but not in samples obtained from *Pcdh19* knockout (KO) mice (Fig. 1C), which instead displayed an additional band of ∼30 kDa, probably reflecting residual expression from exons 4-6, which are still present in this KO model. Interestingly, the CTF1 and 2 bands were much stronger in postnatal than in embryonic lysates (Fig. 1C), indicating that processing is a more common event in neurons than in progenitors, which are the predominant cell type in E11 forebrain. This difference could be due, at least in part, to the reported activity-dependent processing of PCDH19 (Gerosa et al., 2022), which we confirmed by treatment of mESC-derived neurons with N-methyl D-aspartate (NMDA) (Fig. 1D) that led to a significant increase in the intensity of the CTF2 band. Pre-treatment with MK-801, a specific inhibitor of the NMDA receptor, abolished this increase, confirming that activation of the NMDA receptor is sufficient to trigger PCDH19 processing in cortical-like neurons (Ctrl: 0.004 ± 0; 30’ NMDA: 0.33 ± 0.05; 30’ NMDA + MK-801: 0.02 ± 0.02; n = 3; one way ANOVA, F(2,6) = 37.65, *p* = 0.0004; Fig. 1D).

Our data thus indicate that PCDH19 proteolytic processing is a common event, taking place in HEK293 cells, in cortical-like mESC-derived neurons and, more importantly, in the cortex *in vivo.* Moreover, in neuronal cells, the processing is activity dependent, with an increase in processing in response to NMDA receptor activation.

### Importins mediate the nuclear translocation of PCDH19-ICD

The intracellular domain (ICD) of PCDH19 contains a putative nuclear localization signal (NLS) (Fig. S1A) and has been shown to translocate to the nucleus (Gerosa et al., 2022), but the mechanism behind its nuclear import has not been investigated. We confirmed nuclear distribution of PCDH19-ICD in cortical neurons both *in vitro* (Fig. S1D,E) and *in vivo* (Fig. S1F) and further hypothesized that nuclear import of PCDH19-ICD might be mediated by the classical pathway involving importin dimers (Lu et al., 2021). To test whether PCDH19 interacts with importins, we carried out co-immunoprecipitation experiments with PCDH19 and importin subunit alpha 5 (KPNA1) (Fig. 2A). The interaction with PCDH19-FL-HA was weaker than with PCDH19-ICD-HA, suggesting that anchorage to the plasma membrane decreases the probability of interaction between the two proteins. We also verified the interaction between PCDH19 and KPNA1 in the cortex *in vivo*, by co-immunoprecipitation using P10 cortical lysates (Fig. 2B). Finally, to verify that this interaction is mediated by the predicted bipartite NLS (aa 762 to 780: “KRIAEYSYGHQKKSSKKKK”) and that the NLS is functional, we mutated its last four basic residues from “KKKK” to “AAAA” in the PCDH19-ICD-HA construct. As expected, expression of this construct in HeLa cells resulted in a significant reduction in nuclear signal ratio compared to non-mutated PCDH19-ICD-HA (19ICD: 82.17 ± 2.84; 19ICD_(NLSmut)_: 53.62 ± 3.51, n = 15; Mann-Whitney, *p* < 0.0001; Fig. 2C). The retained nuclear signal could be explained by a secondary import mechanism, as PCDH19 also contains two predicted monopartite NLS (aa 1031 to 1037: “PTLKGKR” and aa 1138 to 1144 “PGVKRLK”). Therefore, our data show that processing of PCDH19 generates a soluble cytoplasmic fragment with a functional NLS that is recognized by importins for nuclear transport.

**Figure 2.**
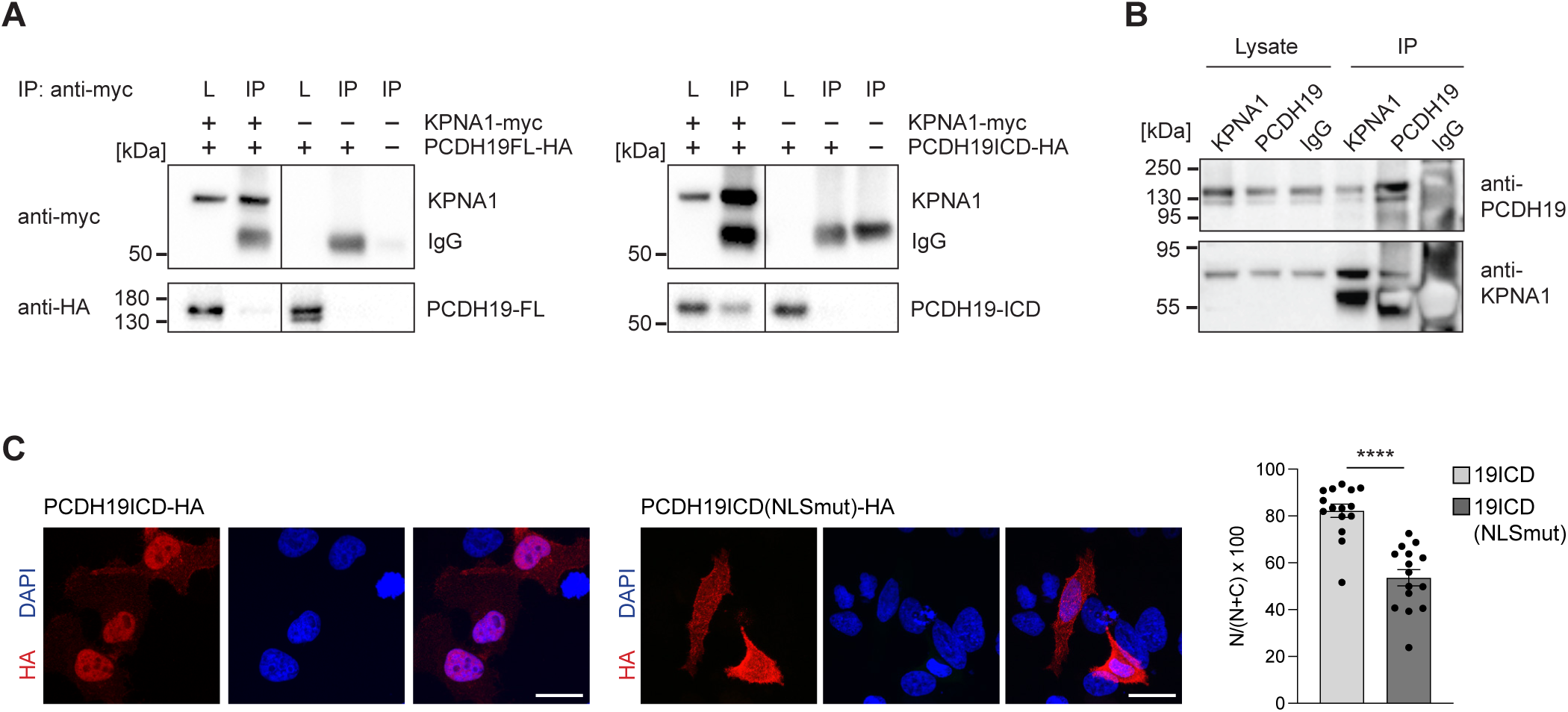
Importins mediate PCDH19-ICD nuclear transport. A. Co-IPs of PCDH19FL (left) and PCDH19-ICD (right) with importin α1 (KPNA1). B. Co-IP of PCDH19 and KPNA1 from P10 cortical lysates. IPs were performed with anti-PCDH19, anti-KPNA1 and IgG. C. HeLa cells transfected with PCDH19ICD-HA or PCDH19ICD-HA with a mutated NLS, and subjected to a 20 min ionomycin treatment. Representative confocal images of transfected cells immunostained for HA (left). Nuclei are counterstained with DAPI. Relative nuclear fluorescence was quantified, mean ± SEM is displayed (right, N=15). Scale bar: 20 μm. Data information: Relative nuclear fluorescence was obtained by dividing nuclear fluorescence by the sum of nuclear and cytoplasmic fluorescence. N: Individual cells. ****p < 0.0001. Statistical analysis: unpaired two-tailed Student’s t-test.

### Constitutive expression of PCDH19 ICD alters the expression of genes involved in neuronal differentiation and synaptic function

Our results indicate that proteolytic processing of PCDH19 produces a soluble ICD fragment that can enter the nucleus, suggesting a potential role in gene regulatory function for this protein. However, although PCDH19 is known to regulate expression of immediate early genes (Gerosa et al., 2022), it is still unknown what impact PCDH19-ICD has on the transcriptional landscape of neurons. To address this question and to examine the potential gene regulatory function of PCDH19-ICD in neuronal cells, we generated a transgenic mESC line with constitutive expression of an HA-tagged cytoplasmic domain from the *Rosa26* locus in addition to the endogenous gene (Fig. 3A and Fig. S2). Bulk RNAseq analysis was then performed with samples obtained from three independent differentiations of control and 19ICD-OE mESC-derived neurons at day *in vitro 8* (DIV8) and DIV12.

**Figure 3.**
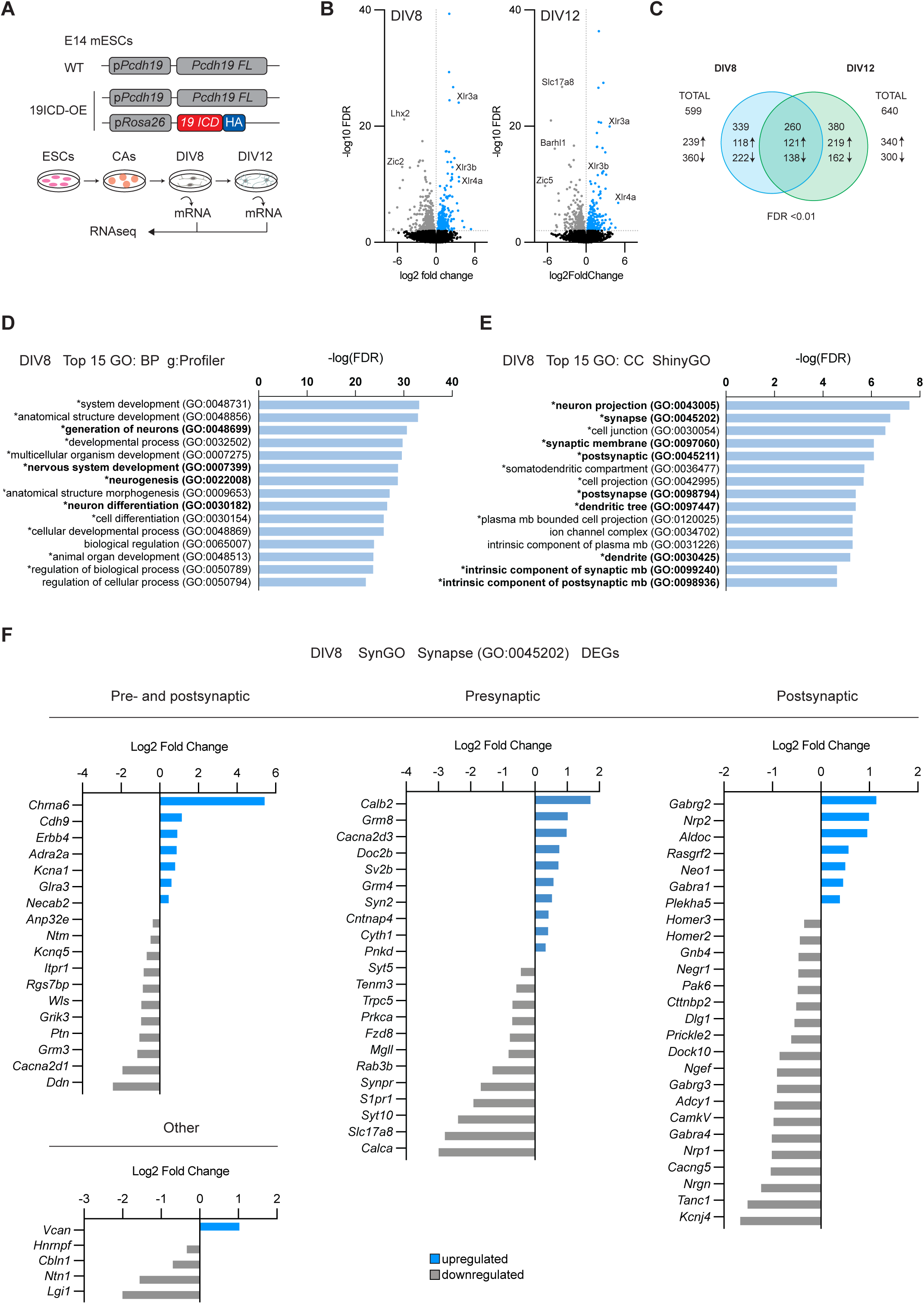
Overexpression of PCDH19-ICD leads to transcriptional changes in neuronal differentiation and synaptic genes in mESC-derived neurons. A. Experimental strategy. B. Volcano plots of DEGs at DIV8 and DIV12. C. Venn diagram with total numbers of DEGs and comparison of common and unique up- and downregulated genes between WT (N=3) and 19ICD-OE (N=3) at DIV8 and DIV12, determined by RNAseq. The number of up- and downregulated genes is indicated by upward and downward arrows, respectively. D. Top 15 Biological Process GO-terms enriched among the differentially expressed genes at DIV8 as determined by gProfiler. Log(FDR) values are shown for each term. Common top 15 BP GO-terms between gProfiler, ShinyGO and Panther are marked with an asterisk and terms relevant to neuronal differentiation are highlighted in bold. E. Top 15 Cellular Compartment GO-terms enriched among the differentially expressed genes at DIV8 as determined by ShinyGO. Log(FDR) values are shown for each term. Common top 15 CC GO-terms between gProfiler, ShinyGO and Panther are marked with an asterisk and terms related to neuronal dendrites and synapses are highlighted in bold. F. List of significantly up- and downregulated genes between DIV8 WT and 19ICD-OE mESC-derived neurons related to the GO-term “Synapse” as determined by RNAseq. Genes are classified according to their synaptic location, and their log2 fold change is depicted in blue (upregulated) or grey (downregulated).

To investigate specific effects of PCDH19-ICD overexpression on the neuronal transcriptional landscape, we carried out differential expression analysis at each time point (DIV8 and DIV12), comparing 19ICD-OE and WT neurons. Analysis with a false discovery rate (FDR) cutoff of 0.01 identified 599 differentially expressed genes (DEGs) at DIV8 and 640 DEGs at DIV12 (Fig. 3B). A comparison between DIV8 and DIV12 DEGs revealed 260 common genes, representing 43% of DIV8 and 41% of DIV12 DEGs, respectively. In all cases except one (the predicted lincRNA *Gm26586*), the direction of change was maintained, suggesting continuous regulation of those genes by PCDH19-ICD over the course of neuronal maturation (Fig. 3C). Verification of differential expression by real time PCR was performed for several genes (*Zic1*, *Lhx2* and *Erbb4*) (Fig. S2).

Insight into the types of genes and processes affected by the transcriptional changes was obtained by subjecting the identified DEGs to an over-representation analysis matching against gene ontology (GO) terms. We compared the results obtained with g:Profiler (Raudvere et al., 2019), ShinyGO (Ge et al., 2019) and Panther (Mi et al., 2010; Thomas et al., 2003) and provide the average FDR value (FDR_A_) for each term. Thirteen of the top 15 GO terms for biological function at DIV8 were shared between all three platforms, with four of them directly related to neural development: “nervous system development” (FDR_A_= 1.11E-25), “neurogenesis” (FDR_A_= 1.50E-23), “generation of neurons” (FDR_A_ = 5.99E-24) and “neuron differentiation” (FDR_A_ = 6.75E-21) (Fig. 3D). The remaining GO terms, whether shared or not, were all related to developmental and differentiation processes (Fig. 3D). At DIV12, these four terms were still significant across the three platforms, and they still ranked among the top 15 in Panther and ShinyGO, with broader terms related to differentiation and transcription ranking higher in gProfiler.

In the cellular component (CC) category, 13 of the top 15 GO terms were again shared between the three platforms at DIV8, including terms such as “neuron projection” (FDR_A_ = 9.12E-09), “cell junction (FDR_A_ = 8.93E-08) or “synapse” (FDR_A_ = 5.58E-08) and many of the significant terms were directly or indirectly related to dendrites and synapses (Fig. 3E). Therefore, we run our list of DEGs through the SynGO database, which highlighted several pre- and postsynaptic genes, including postsynaptic scaffolding proteins, neurotransmitter receptors, synaptic vesicle proteins, channels and channel subunits, cell adhesion proteins and others (Fig. 3F). Interestingly, about two thirds of the synaptic DEGs were downregulated (Fig. 3F). The biological processes “trans-synaptic signalling” (FRD = 0.0019), “chemical synaptic transmission” (FRD = 0.0048) and “presynaptic modulation of chemical synaptic transmission” (FRD = 0.0019) were also significant in this database.

Ingenuity pathway analysis (IPA) of diseases and functions of the differentially expressed genes produced similar results, with the function “differentiation of neurons” showing the lowest P-value at DIV8 (P_(DIV8)_ = 6.4E-20) and the ninth lowest at DIV12 (P_(DIV12)_ = 4.03E-13). Other related functions, such as “development of neurons” (P_(DIV8)_ = 1.61E-09; P_(DIV12)_ = 8.42E-06), “neuritogenesis” (P_(DIV8)_ = 2.74E-06; P_(DIV12)_ = 1.07E-04) and “guidance of axons” (P_(DIV8)_ = 8.12E-09; P_(DIV12)_ = 9.54E-07) also displayed highly significant P values at DIV8 and DIV12. Interestingly, the z-score associated with the pathway “differentiation of neurons” reached the threshold to be predicted as decreased at DIV8 (z-score −2.318), although not at DIV12 (z-score −1.234). Furthermore, two functions related to synaptic function, “synaptic transmission” (P_(DIV8)_ = 6.31E-06; z-score −2.158) and “neurotransmission” (P_(DIV8)_ = 3.65E-05; z-score −2.359) were also predicted to be decreased at DIV8, indicating that PCHD19-ICD might negatively regulate some of those processes, at least during early stages, or that compensatory mechanisms emerge as development progresses.

Together, our results show that PCDH19-ICD broadly affects the neuronal transcriptome and, in particular, the expression of genes involved in processes of neuronal differentiation, maturation and synaptic transmission.

### Expression of PCDH19-ICD in cortical neurons reduces the number of spines without affecting neuronal morphology

Considering the results obtained in our transcriptomic analysis, with DEGs linked to GO terms related to different processes of neuronal differentiation and synaptic function, we decided to investigate the role of PCDH19-ICD *in vivo*, to better understand the roles that proteolytic processing of this cell adhesion protein plays in the cortex. Because the processing sites for PCDH19 are currently unknown, and the proteases involved seem to differ between different neuronal types, it was not possible to pursue a loss of function approach in which proteolytic processing of PCDH19 is prevented. Therefore, we opted for an overexpression paradigm and carried out functional analyses using *in utero* electroporation at embryonic day E15.5 to target cortical layer 2/3, one of the main cortical populations known to express *Pcdh19* (Galindo-Riera et al., 2021). We electroporated control plasmid pCIG or pCIG-PCDH19-ICD-HA (abbreviated pCIG-19ICD-HA) and analysed mature neurons at postnatal day P60 after immunostaining with anti-HA and anti-EGFP antibodies (Fig. 4A,B). After ruling out any effects of PCDH19-ICD on neurogenesis, radial migration and axonal guidance (Fig. S3), we checked if the changes to the transcriptional landscape elicited by this fragment had an impact on overall neuronal morphology. To this end, we imaged and traced electroporated neurons to evaluate any potential changes in dendritic arborization (Fig. 4C,D). No differences were found in the width of the apical or basal dendritic arbors (apical, pCIG: 284.55 ± 33.99 μm, n = 11; pCIG-19ICD-HA: 266.29 ± 21.79 μm, n = 12; Mann-Whitney, *p* = 0.9279; basal, pCIG: 273.23 ± 25.74 μm, n = 11; pCIG-19ICD-HA: 261.59 ± 11.53 μm, n = 12; Mann-Whitney, *p* = 0.9759; Fig. 4E) nor in the total added length of dendrites of the different orders (order 1, pCIG: 635.63 ± 50.62 μm, n = 11; pCIG-19ICD-HA: 576.2 ± 88.5 μm, n = 12; independent t-test, *p* = 0.5755; order 2, pCIG: 1294.99 ± 106.45 μm, n = 11; pCIG-19ICD-HA: 1198.27 ± 107.49 μm, n = 12; independent t-test, *p* = 0.5305; order 3, pCIG: 1598.32 ± 185.51 μm, n = 11; pCIG-19ICD-HA: 1473.14 ± 210.04 μm, n = 12; independent t-test, *p* = 0.6621; order 4, pCIG: 667.7 ± 146.87 μm, n = 10; pCIG-19ICD-HA: 606.9 ± 116.48 μm, n = 11; independent t-test, *p* = 0.747; order 5, pCIG: 153.27 ± 50.88 μm, n = 7; pCIG-19ICD-HA: 287.68 ± 108.69 μm, n = 9, Mann-Whitney, *p* = 0.4874; Path lengths 6 to 8 could not be tested due to low numbers; Fig. 4F). Sholl analysis didn’t show any other morphological differences between neurons electroporated with pCIG or with pCIG-19ICD-HA (Fig. 4G). These results suggest that the transcriptional changes elicited by PCDH19-ICD either do not directly affect neuronal morphology *in vivo*, or can be compensated during development, giving rise to morphologically unaltered neurons.

**Figure 4.**
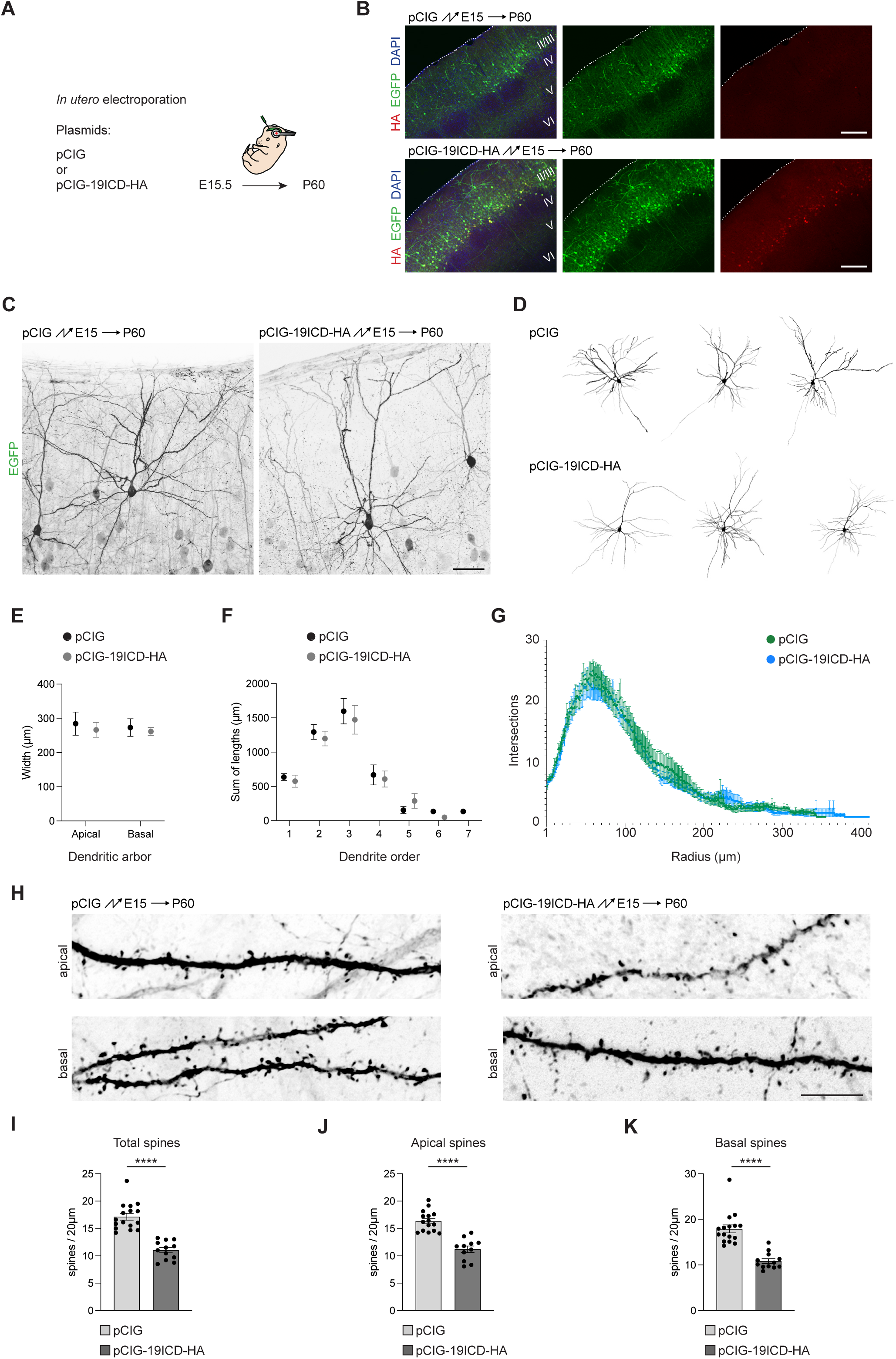
*In vivo* overexpression of PCDH19 ICD reduces spine density without affecting dendritic morphology. A. Strategy for the *in utero* electroporation to express PCDH19 ICD in upper layer cortical neurons. B. Representative confocal images of electroporated brains immunostained for HA for the control (pCIG) and experimental (pCIG-19ICD-HA) conditions. Cortical layers are indicated with roman numerals. Scale bar: 200 μm. C. Maximum intensity projections of confocal stacks showing individual upper layer neurons targeted with pCIG (left) or pCIG-19ICD-HA (right). Fluorescent images have been inverted and are shown in greyscale to improve contrast. Scale bar: 50 μm. D. Examples of traced neurons electroporated with pCIG (top) or pCIG-19ICD-HA (bottom). E. Quantification (mean ± SEM) of the width of the apical and basal dendritic arbors of pCIG (N=11 neurons, from N=6 animals) or pCIG-19ICD-HA (N=12 neurons, from N=6 animals) electroporated neurons, determined with the SNT plug in in Fiji. F. Quantification (mean ± SEM) of the total dendritic length by dendrite order in pCIG (N=11 neurons, from N=6 animals) or pCIG-19ICD-HA (N=12 neurons, from N=6 animals) electroporated neurons, determined with the SNT plug in in Fiji. G. Sholl analysis of pCIG (N=11 neurons, from N=6 animals) and pCIG-19ICD-HA (N=12 neurons, from N=6 animals) electroporated neurons, determined with the SNT plug in in Fiji. H. Maximum intensity projections of confocal stacks showing apical and basal dendrites of pCIG and pCIG-19ICD-HA electroporated neurons at high magnification to visualize dendritic spines (63x objective). Fluorescent images have been inverted and are shown in greyscale to improve contrast. Scale bar: 10 μm. I-K. Quantification (mean ± SEM) of total (I), apical (J) and basal (K) spine density in second order dendrites of pCIG (N=16) and pCIG-19ICD-HA (N=12) electroporated neurons. In each case, those neurons came from 6 different electroporated animals. Data information: (E,F) N: Individual neuron. No statistically significant differences identified. Statistical analysis: two-tailed Mann-Whitney test for apical and basal dendritic arbors (E); unpaired two-tailed Student’s t-test for orders 1-4, two-tailed Mann-Whitney test for dendrite order 5. Analysis of orders 6 and 7 was not possible due to low numbers (F). (I-K): The average of all dendritic areas analysed per neuron was used for the analysis of spine density. N: Individual neuron. Relevant p-values are indicated: ****p < 0.0001. Statistical analysis: two-tailed Mann-Whitney test for total and basal spine densities (I, K); unpaired two-tailed Student’s t-test for apical spine density (J).

PCDH19 localizes at synapses and its proteolytic processing is activity dependent, pointing to a potential synapse to nucleus signalling pathway. In addition, the GO term analysis of our RNAseq experiment returned several significant synaptic terms both in the biological function and cellular compartment categories (Fig. 3D-F), further suggesting a role for PCDH19-ICD in synaptic signalling. In that analysis, many postsynaptic genes were downregulated, and IPA analysis also predicted synaptic transmission to be reduced. Therefore, we hypothesized that if PCDH19-ICD is affecting synaptic transmission, it could directly or indirectly impact differentiation processes related to spine formation or stabilization, since the main morphological manifestation of excitatory synapses are dendritic spines. To test this hypothesis, we quantified the number of spines in dendrites from pCIG and pCIG-19ICD-HA electroporated neurons and calculated the spine density. Interestingly, pCIG-19ICD-HA electroporated neurons showed a ∼36% reduction in total spine density (pCIG: 17.16 ± 0.62 spines/20μm, n = 16; pCIG-19ICD-HA: 11.04 ± 0.49 spines/20μm, n = 12; Mann-Whitney, *p* < 0.0001, Fig. 4H,I). This reduction was slightly more pronounced across basal (∼40%) than apical (∼32%) spines (apical, pCIG: 16.36 ± 0.49 spines/20μm, n = 15; pCIG-19ICD-HA: 11.20 ± 0.61 spines/20μm, n = 11; independent t-test, *p* = 6.75E-07; Fig. 5J; basal, pCIG: 17.91 ± 0.87 spines/20μm, n = 16; pCIG-19ICD-HA: 10.87 ± 0.53 spines/20μm, n = 12; Mann-Whitney, *p* < 0.0001; Fig. 5K).

**Figure 5.**
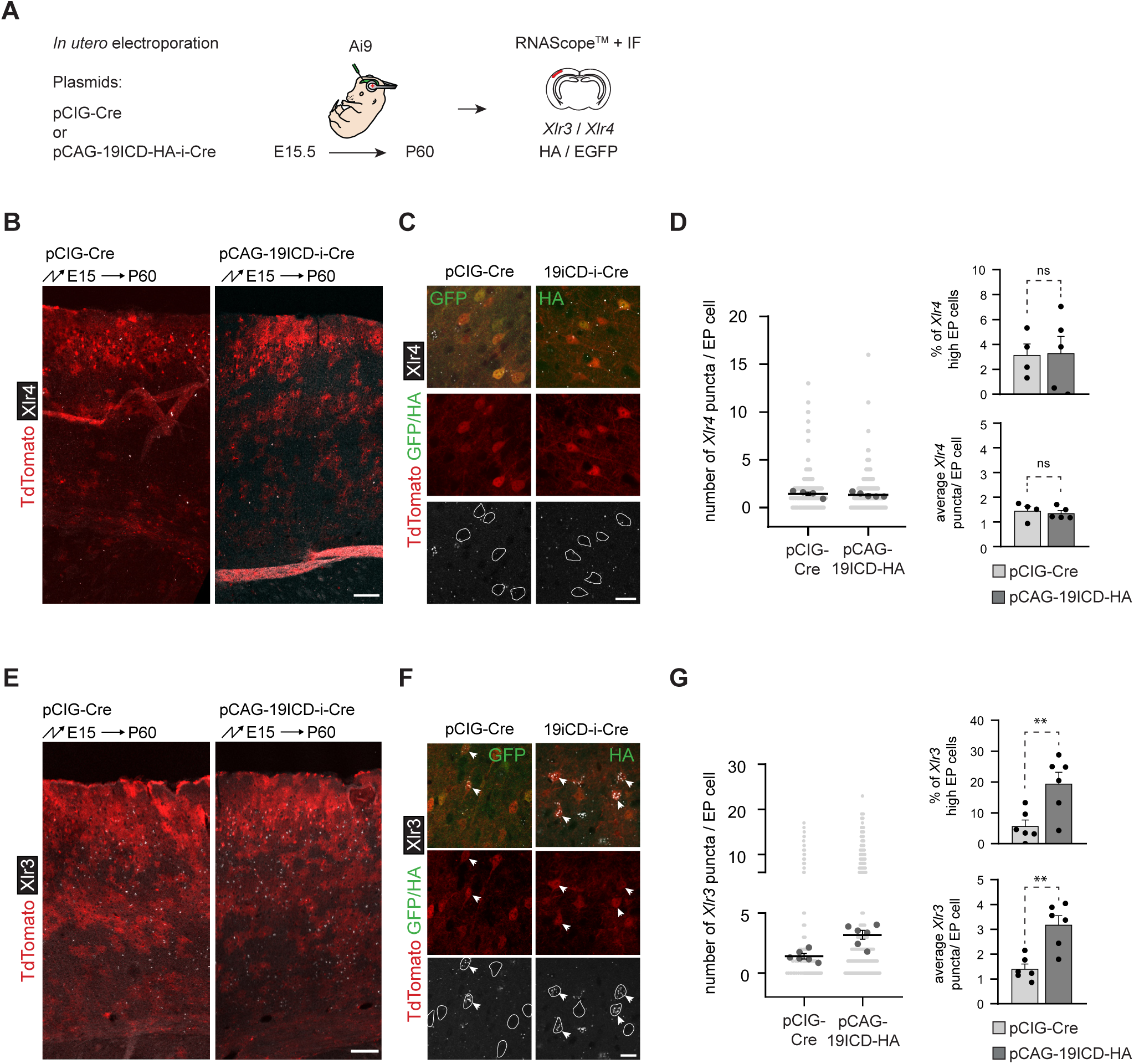
Electroporation of PCDH19 ICD leads to overexpression of *Xlr3* genes. A. Experimental strategy. B. General overview of pCIG-Cre and pCAG-19ICD-HA-ires-Cre electroporated hemispheres after *Xlr4* RNAScope. Scale bar: 500 μm. C. Representative confocal images of pCIG-Cre and pCAG-19ICD-HA-ires-Cre electroporated neurons displaying EGFP (left) or HA (right) immunofluorescence and RNAScope signal for *Xlr4* genes. Cell bodies of electroporated neurons are highlighted. Scale bar: 20 μm. D. Quantification of the number of *Xlr4* puncta per cell for pCIG-Cre (N=4, from 4 different litters) and pCAG-19ICD-HA-ires-Cre (N=5, from 4 different litters) electroporated brains. Data from all electroporated cells counted (light grey dots), average per animal (dark grey dots) and average per condition (black line) are shown on the left, average percentage of ‘*Xlr4* high” electroporated cells (top) and average *Xlr4* puncta per electroporated cell (bottom) are shown on the right. E. General overview of pCIG-Cre and pCAG-19ICD-HA-ires-Cre electroporated hemispheres after *Xlr3* RNAScope. F. Representative confocal images of pCIG-Cre and pCAG-19ICD-HA-ires-Cre electroporated neurons displaying EGFP (left) or HA (right) immunofluorescence and RNAScope signal for *Xlr3* genes. Cell bodies of electroporated neurons are highlighted. Arrows point to *Xlr3* high expressing cells. Scale bar: 20 μm. G. Quantification of the number of *Xlr3* puncta per cell for pCIG-Cre (N=6, from 5 different litters) and pCAG-19ICD-HA-ires-Cre (N=6, from 4 different litters) electroporated brains. Data from all electroporated cells counted (light grey dots), average per animal (dark grey dots) and average per condition (black line) are shown on the left, average percentage of ‘*Xlr3* high” electroporated cells (top) and average *Xlr3* puncta per electroporated cell (bottom) are shown on the right. Data information: (D, G) N: Individual brain. Relevant p-values are indicated: **p < 0.01. Statistical analysis: unpaired two-tailed Student’s t-test.

Our results demonstrate that upper layer cortical neurons overexpressing PCDH19-ICD *in vivo* have fewer spines on their dendrites, pointing to a role of this fragment, and thus of the proteolytic processing of PCDH19, in the regulation of spine density.

### PCDH19-ICD increases expression of *Xlr* genes *in vivo*

To better understand how PCDH19-ICD might regulate dendritic spine density, we went back to our RNAseq results to search for genes with a known involvement in spine regulation. We considered the upregulation of several genes of the *Xlr* family in the 19ICD-OE samples particularly interesting, as overexpression of *Xlr* genes has been linked to a decrease in spine density in cortical neurons of the upper layers of the cortex (Cubelos et al., 2010). In our samples, upregulated *Xlr* genes included *Xlr3a* (log_2_ FC: 3.66, FDR: 1.15E-20), *Xlr3b* (log_2_ FC: 2.52, FDR: 1.38E-12) and *Xlr4a* (log_2_ FC: 4.99, FDR: 1.58E-07) (Fig. 3B). *Xlr4b* was also upregulated (log_2_ FC: 4.23), but its very low expression in control samples precluded the calculation of the adjusted p value. We therefore sought to confirm the upregulation of *Xlr* genes in cortical neurons overexpressing PCDH19-ICD *in vivo*, which could underpin the observed spine phenotype. To this end, two RNAScope probes were designed to detect *Xlr3* and *Xlr4* genes, respectively, and used on electroporated brain slices to quantify the expression of *Xlr3* and *Xlr4* genes in neurons overexpressing PCDH19-ICD vs control electroporated neurons (Fig. 5A). Expression of *Xlr4* genes was very low, and no significant differences were found in the average number of transcripts per cell, quantified as the number of fluorescent puncta, nor in the percentage of cells with 5 or more puncta (“*Xlr4* high” cells) (pCIG-Cre: 1.45 ± 0.18 transcripts/cell, n = 4; pCIG-19ICD-HA-i-Cre: 1.35 ± 0.11 transcripts/cell, n = 5; independent t-test, *p* = 0.63; pCIG-Cre: 3.12 ± 0.89%, n = 4; pCIG-19ICD-HA-i-Cre: 3.31 ± 1.35%, n = 5; independent t-test, *p* = 0.44; Fig. 5B-D). Expression of *Xlr3* genes was higher than that of *Xlr4*, and it was affected by PCDH19-ICD overexpression: both the average number of *Xlr3* transcripts per cell and the percentage of “*Xlr3* high” cells, defined in this case as neurons with 6 or more puncta (due to the overall higher expression), were increased in the PCDH19-ICD overexpressing population (*Xlr3*: pCIG-Cre: 1.41 ± 0.19 transcripts/cell, n = 6; pCIG-19ICD-HA-i-Cre: 3.18 ± 0.36 transcripts/cell, n = 6; independent t-test, *p* = 0.0015; pCIG-Cre: 5.68 ± 1.98 %, n = 6; pCIG-19ICD-HA-i-Cre: 19.44 ± 3.73%, n = 6; independent t-test, *p* = 0.0086; Fig. 5E-G). Thus, the overexpression of *Xlr3* genes in the 19ICD-OE samples from the *in vitro* RNAseq analysis is replicated *in vivo*, providing mechanistic insight into the role of PCDH19-ICD in cortical neurons.

### Downregulation of *Xlr* genes restores spine density in PCDH19-ICD overexpressing neurons

Elevated *Xlr* gene expression has previously been linked to diminished spine density in upper layer cortical neurons without affecting overall neuronal morphology (Cubelos et al., 2010), the same phenotype we see when we express PCDH19-ICD in that neuronal type. Since PCDH19-ICD increases expression of *Xlr* genes, we reasoned that the downregulation of this gene family might rescue the spine deficit we see in electroporated neurons. To test this hypothesis, we co-electroporated pCIG-19ICD-HA with *Xlr* shRNAs that have previously been shown to reduce their expression (Cubelos et al., 2010) or with a control shRNA (Fig. 6A). Downregulation of *Xlr3* and *4* genes rescued the spine deficit in neurons overexpressing PCDH19-ICD, increasing spine density by about 30%, while downregulation of *Xlr4* genes alone failed to achieve a significant increase in spine density (pCBA + shCtrl: 26.32 ± 1.87 spines/20μm; pCIG-19ICD-HA + shCtrl: 18.30 ± 0.77 spines/20μm; pCIG-19ICD-HA + shRNA*Xlr3*+*4*: 22.92 ± 1.14 spines/20μm; pCIG-19ICD-HA + shRNA*Xlr4*: 21.31 ± 0.73 spines/20μm, one way ANOVA, F(3,47) = 9.367, *p* < 0.0001; post-hoc Tukey: P < 0.0001 pCBA + shCtrl vs pCIG-19ICD-HA + shCtrl; P = 0.0152 pCIG-19ICD-HA + shCtrl vs pCIG-19ICD-HA + shRNA*Xlr3*+*4* and P = 0.0219 pCBA + shCtrl vs pCIG-19ICD-HA + sh*Xlr4*) (Fig. 6B,C). The fact that reducing *Xlr4* gene expression alone did not rescue the spine phenotype is in accordance with the RNAScope data showing that *Xlr4* genes are not upregulated by PCDH19-ICD expression in cortical neurons *in vivo*. Our findings thus suggest that the observed reduction in spine density upon PCDH19-ICD expression is mediated by the upregulation of *Xlr3* genes.

**Figure 6.**
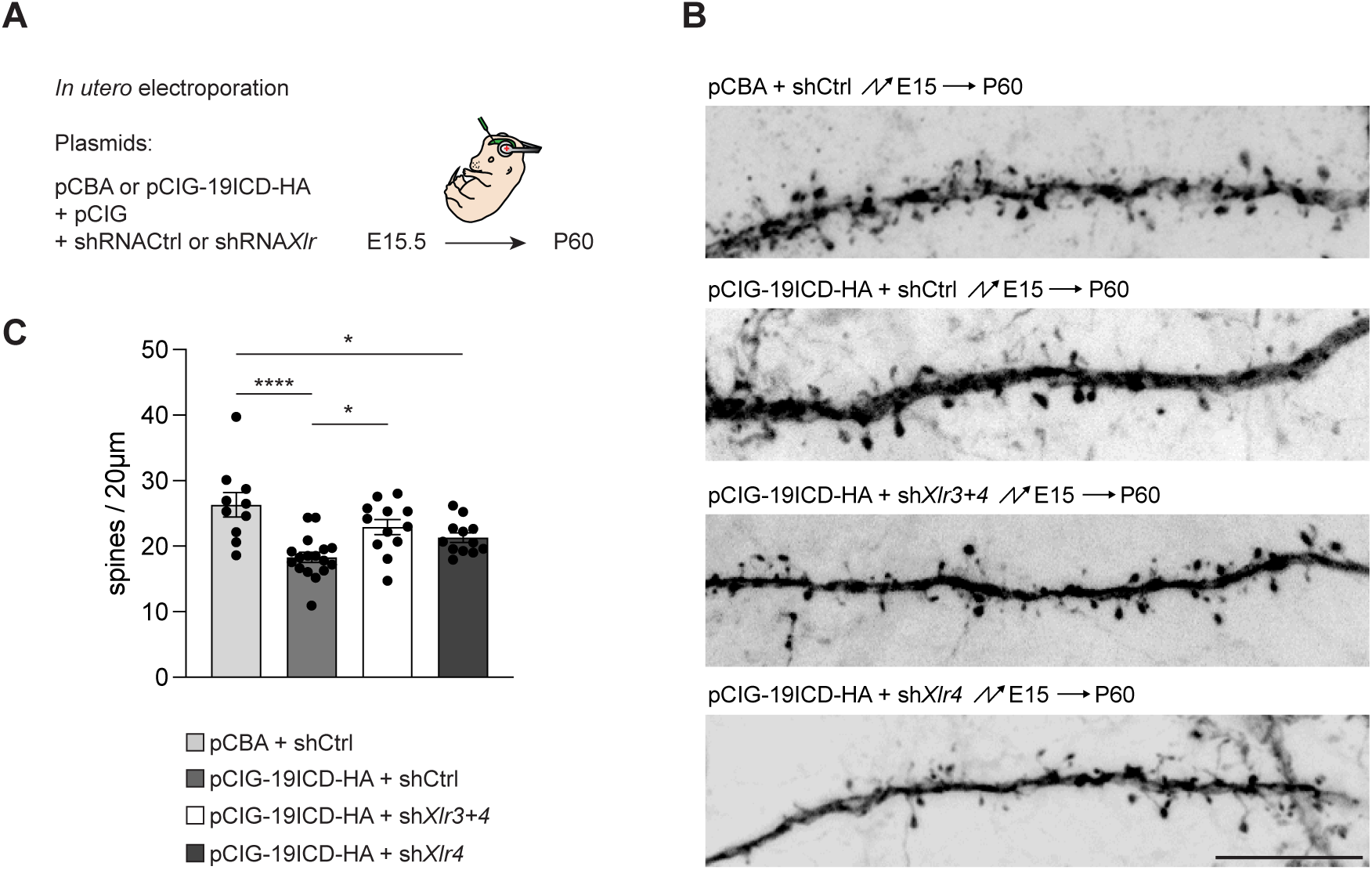
Downregulation of *Xlr* genes rescues spine density in neurons electroporated with PCDH19 ICD. A. Experimental strategy. B. Maximum intensity projections of confocal stacks showing dendrites of neurons electroporated with either pCBA + shCtrl, pCIG-19ICD-HA + shCtrl, pCIG-19ICD-HA + shXl*r3+4* or pCIG-19ICD-HA + sh*Xlr4,* at high magnification to visualize dendritic spines (63x objective). Fluorescent images have been inverted and are shown in greyscale to improve contrast. Scale bar: 10 μm. C. Quantification (mean ± SEM) of total spine density in second and third order dendrites of pCBA + shCtrl (N=10), pCIG-19ICD-HA + shCtrl (N=17), pCIG-19ICD-HA + shXl*r3+4* (N=12) or pCIG-19ICD-HA + sh*Xlr4* (N=12) electroporated neurons. In each case, neurons came from 3-4 different electroporated animals from 2-3 different litters. Data information: (C) The average of all dendritic areas analysed per neuron was used for the analysis of spine density. N: Individual neuron. Relevant p-values are indicated: *p < 0.05, ****p < 0.0001. Statistical analysis: one-way ANOVA with Tukey’s post-hoc test.

## DISCUSSION

PCDH19 is a cell adhesion protein mutated in a form of epileptic encephalopathy (Dibbens et al., 2008), but the way in which it contributes to the disorder is complex and currently not understood. We have investigated here non-adhesive functions of PCDH19, and we provide evidence that its intracellular domain regulates expression of genes involved in neuronal differentiation and controls spine density in upper layer cortical excitatory neurons. Our biochemical experiments demonstrate that proteolytic cleavage of PCDH19, which is activity dependent in neurons, generates a soluble fragment that interacts with importins to be transported into the nucleus. Using RNAseq, we further show that the intracellular domain of PCDH19 elicits broad transcriptional changes associated with synapses and neuronal differentiation processes in mESC-derived neurons. Our *in vivo* functional analysis indicates that overexpression of PCDH19-ICD in layer 2/3 cortical neurons leads to a reduction in apical and basal spine density via upregulation of *Xlr3* genes. Our data thus point to a new way in which PCDH19 might contribute to the pathophysiology of the epileptic encephalopathy it is associated with.

Proteolytic processing has been described for members of different cell adhesion families, including neuroligins, NCAM and cadherins (Lee and Ch’ng, 2020). Processed cadherins include CDH1, CDH2, PCDH12 and the clustered alpha and γ-protocadherins, and in all cases ADAM10 and γ-secretase play a role in the proteolytic steps (Reiss et al., 2005; Maretzky et al., 2005; Reiss et al., 2006; Bonn et al., 2007; Haas et al., 2005; Marambaud et al., 2003, 2002; Uemura et al., 2006; Hambsch et al., 2005; Bouillot et al., 2011). A similar scenario has recently been described for PCDH19 in cultured hippocampal neurons (Gerosa et al., 2022), again involving ADAM10 and, possibly, γ-secretase. Our results confirm the proteolytic cleavage of PCDH19 and its activity dependence in neuronal cells but seem to rule out a major sheddase role for ADAM10 in cortical-like neurons. Instead, other metalloproteases or ADAMs are likely to be involved in these cells. Thus, the specific mechanism of PCDH19 processing might be cell-type dependent, or different proteases might be triggered to cut by specific stimuli depending on the cellular context.

Soluble fragments generated by proteolytic processing of membrane proteins can display further biological functions inside the cell. Interestingly, nuclear translocation of the processed fragments has been described for several CAMs, like ERBB4 (Allison et al., 2011), L1 (Minguez et al., 2020) or DSCAM/DSCAML (Sachse et al., 2019). Within the cadherin superfamily, the ICDs of CDH2 and the γ-protocadherins also localize to the nucleus, and the nuclear functions of these fragments have been characterized to some extent. CDH2-ICD binds to CBP, leading to its degradation and a reduction in CREB dependent transcriptional activity (Marambaud et al., 2003), while γ-protocadherin ICD leads to an increase in γ-protocadherin expression, indicative of an autoregulatory loop (Hambsch et al., 2005). However, genome wide analyses of the changes that nuclear ICDs elicit at the transcriptional level are still scarce and include DSCAM and DSCAML in HEK293 cells (Sachse et al., 2019), and ERBB4 using microarrays in rat cultured hippocampal neurons (Allison et al., 2011), but no such data are available for members of the cadherin superfamily. We have evaluated here the alterations in the transcriptional landscape of mESC-derived neurons constitutively expressing PCDH19-ICD and have found broad changes in gene expression programs relevant for neuronal differentiation. Given the expression profile of *Pcdh19*, which peaks at about postnatal day 7 in the mouse cortex and is then maintained throughout adulthood, and the fact that proteolytic processing of PCDH19 is activity dependent, our results point to a role of this protein in the regulation of neuronal differentiation in response to neuronal activity.

Assessing the *in vivo* relevance of CAM proteolytic events is challenging, not least because of the lack of conserved protease recognition sequences that would facilitate the generation of cleavage defective versions. Only recently has a CDH2 cleavage resistant mouse model been characterized (Asada-Utsugi et al., 2021) that displays spine and synapse anomalies in the hippocampus and enhanced spatial memory. We acknowledge the lack of a loss of function approach in this study. However, unlike for CDH2 (Uemura et al., 2006), the specific sites at which PCDH19 is processed are currently unknown. Furthermore, our data show that PCDH19 might be processed by different proteases depending on cell or even neuronal type, making such an approach currently not viable.

Impairing nuclear translocation of PCDH19-ICD is not without challenges, either. Different mutations in the described bipartite NLS do not completely abolish nuclear localization (Fig. 2F and (Gerosa et al., 2022)), possibly due to the existence of other signals mediating nuclear transport. Widespread mutation to target all potential signals would therefore risk altering other PCDH19 dependent functions. In addition, a novel mouse model would need to be generated to prevent endogenous PCDH19 from being processed and reaching the nucleus. A loss of function approach with shRNA is also not a viable option because, unlike in the hippocampus, expression of *Pcdh19* is not uniform across cortical neurons within a layer, not even in those layers with strongest *Pcdh19* expression (2/3 and 5). Therefore, it would be impossible to know if the targeted neurons expressed *Pcdh19* in the first place. Furthermore, such a strategy would also alter adhesion, possibly leading to confounding effects. For those reasons, our approach to evaluate the *in vivo* functions of PCDH19-ICD has been the opposite, overexpressing it in ESC-derived cortical-like neurons to assess changes to the transcriptome and in upper layer cortical neurons to evaluate the consequences of excessive processed fragment *in vivo*. This overexpression approach, which also reduces potential compensatory effects, has been used to investigate the role of other processed cell adhesion proteins in the past (Sachse et al., 2019) and has allowed us to identify a role for PCDH19-ICD in the regulation of spine number, with continuous expression leading to a reduction in apical and basal spine density through upregulation of *Xlr* gene levels.

The *Xlr* (X-linked lymphocyte regulated) gene family comprises several genes and pseudogenes located on the X chromosome that were identified originally for their expression in lymphoid cell lines (Cohen et al., 1985). Their functions are still mostly unclear, but they have been potentially linked to chromatin metabolism due to the presence of a Cor1/Xlr/Xmr conserved region in their coding sequence and because XLR1 colocalizes with SATB1 in thymocytes (Escalier et al., 1999).

Interestingly, some *Xlr* genes display genetic imprinting (Davies et al., 2005; Raefski and O’Neill, 2005) and at least *Xlr3b* has been associated with a cognitive phenotype in mice (Davies et al., 2005). More recently, *Xlr3b* and *Xlr4b* were shown to regulate dendritic spines in upper layer cortical neurons, acting downstream of the transcription factors *Cux1* and *Cux2* (Cubelos et al., 2010). Loss of *Cux2* resulted in their upregulation, leading to decreased spine density without other changes in neuronal morphology, the same phenotype we have reported here. *Xlr4* genes have also been linked to cocaine addiction (Segni et al., 2020) and several *Xlr* genes from the Xlr3/4/5 cluster were shown to be upregulated in the hippocampus upon USF-1 deficiency (Sideromenos et al., 2022).

Remarkably, upregulation of *Xlr* genes in hippocampal pyramidal neurons led to a different phenotype of shorter dendrites with no change in dendritic spine density, suggesting that differences in the specific combination of upregulated *Xlr* genes or in the neuronal populations on which they act might affect neuronal morphology in different ways.

To elucidate the pathophysiological underpinnings of the epileptic encephalopathy caused by mutations in PCDH19, it is necessary to understand the different functions of this protein, including the less studied, non-adhesive, nuclear roles as the one described here that might offer new insights into the molecular mechanisms at play in this disorder. Our results suggest an involvement of PCDH19 in synaptic homeostasis, wherein neuronal activity would trigger its proteolytic processing, and the resulting fragment would then regulate synaptic numbers through a nuclear signalling and transcriptional pathway mediated by *Xlr* genes. Because PCDH19 and CDH2 are known to interact (Biswas et al., 2010; Emond et al., 2011), CDH2 is also processed in response to neuronal activity (Marambaud et al., 2003), and cleavage resistant CDH2 leads to increased spine density (Asada-Utsugi et al., 2021), it will be interesting to investigate in the future if and how these two cadherins cooperate to maintain optimal spine density in excitatory neurons. It would also be interesting to investigate the *in vivo* roles of PCDH19-ICD both at earlier stages and after neuronal stimulation to evaluate relative contributions of PCDH19 to spine formation and/or maintenance during development and to spine homeostasis in adulthood. Furthermore, given the spatial distribution of PCDH19 expression in the adult cerebral cortex, which is specific to some layer 2/3 and layer 5 neuronal populations (Galindo-Riera, et al 2021), it will also be necessary to complement our findings with an investigation into the contribution of the PCDH19 intracellular fragment to spine homeostasis in layer 5 cortical neurons.

## Data availability

The complete RNAseq data will be available in the GEO repository.

## Acknowledgements

This work was supported by the PCDH19 Alliance (research grant to I.M-G.), the Life Science Research Network Wales, an initiative funded through the Welsh Government’s Ser Cymru program (initial fellowship to I.M-G.), the Wellcome Trust (Seed Award 109643/Z/15/Z to I.M-G., fellowship 204021/Z/16/A to S.A.N., fellowship 220002/Z/19/Z to I.W.J.F.), the Biotechnology and Biological Sciences Research Council (BBSRC) (grant BB/S002359/1 to I.M-G.), the Ministerio de Ciencia e Innovación (PID2020-114227RB-I00 MICIU/AEI/10.13039/501100011033; CNS2022-135758 MICIU/AEI/10.13039/501100011033 and European Union NextGenerationEU/PRTR; PID2023-153143OB-I00 MICIU/AEI /10.13039/501100011033 and FEDER, EU to C.G.-S) and the Consellería de Educación, Universidades y Empleo (Generalitat Valenciana) (CIDEXG/2022/31 to I.M-G.). C.G-S. received a Ramón y Cajal Grant from the MICINN (RyC-2015-19058). J.F-B and V.B-E are recipients of a predoctoral contract from the Generalitat Valenciana (ACIF/2021/139 and CIACIF/2023/456, respectively). We would like to thank James Wilding for initial experiments that didn’t make it into this manuscript and Rebecca Hughes for her participation in the cloning of PCDH19ICD(NLSmut)-HA and IPO5-myc. We would also like to thank Prof Yves Barde, Dr David Petrik and the members of their labs, as well as Prof Ulrich Mueller for their useful feedback during the completion of this work, and Dr Eva Porlán for her insightful comments on proteolytic processing analysis.

Current address for SAN: Achucarro Basque Center for Neuroscience, Leioa 48940, Bizkaia, Spain. Current address for IWJF: University of Edinburgh, Edinburgh EH16 4UU, Scotland, UK Current address for CLl-B: Centre for Developmental Neurobiology, King’s College London, London SE1 1UL, UK.

## Author contributions

SAN, CG-S and IM-G designed research; SAN, VB-E, JF-B, IWJF, CLl-B, ES, CG-S and IM-G performed research; SAN, VB-E, JF-B, IWJF, CLl-B, ES, CG-S and IM-G analyzed data; IM-G wrote the paper with input from SAN, VB-E, JF-B, IWJF, CLl-B, ES and CG-S.

---

**Figure S1.**
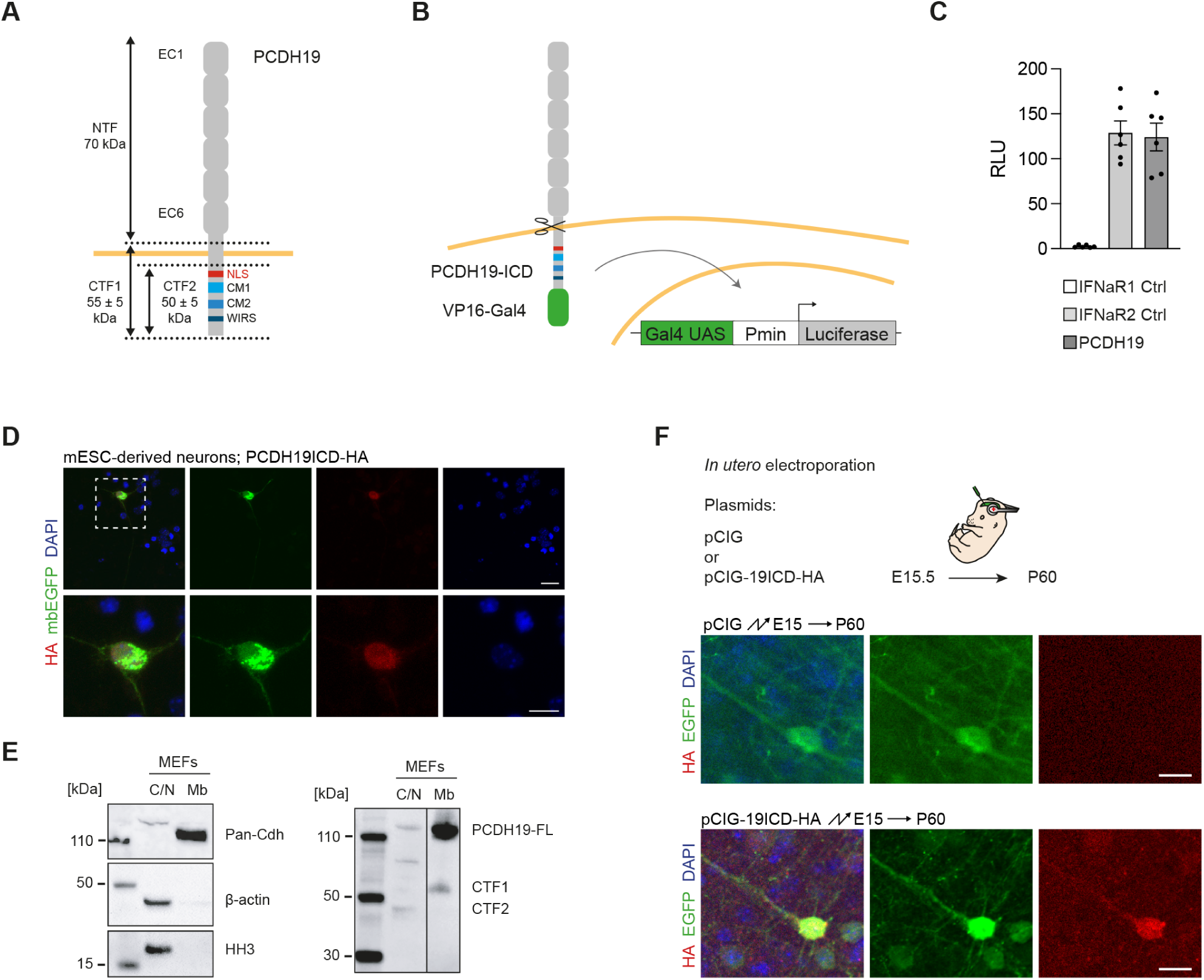
Independent verification of PCDH19 proteolytic processing and nuclear transport of its ICD. A. Schematic of the PCDH19 protein indicating known domains and expected sizes after proteolytic processing. NTF: N-terminal fragment; CTF: C-terminal fragment; NLS: nuclear localization signal; CM: conserved motif; WIRS: WAVE interacting receptor sequence. B. Illustration of the luciferase reporter assay to investigate PCDH19 processing. C. Quantification (mean ± SEM) of the luciferase reporter assay in HEK293 cells. Results are shown as Firefly/Renilla luciferase ratio. IFNaR1 is a negative control and IFNaR2 is a positive control known to be constitutively processed. IFNaR1 Ctrl: 2.56 ± 0.58, IFNaR2 Ctrl: 128.8 ± 13.32, PCDH19-FL: 124.2 ± 15.5; one way ANOVA, F(2,15) = 36.79, *p* < 0.0001. (N=6) D. Representative confocal images of mESC-derived neurons transfected with PCDH19-ICD-HA and immunostained for HA. Nuclei are counterstained with DAPI. Scale bars: 20 μm (overview, top rows), 10 μm (inserts, bottom rows). E. Western blot detection of endogenous full length and processed PCDH19 in cytoplasm/nucleus (C/N) and membrane (Mb) enriched fractions of MEF lysates (right). A verification of the subcellular fractionation of MEF lysates into cytoplasm/nucleus enriched (β-actin, HH3) and membrane enriched (Pan-Cadherin) fractions is shown on the left. F. *In utero* electroporation to express PCDH19-ICD in upper layer cortical neurons. Strategy (top) and representative confocal images of a layer II/III neuron immunostained for HA for the control (pCIG) and 19ICD-HA conditions. Scale bar: 20 μm. Data information: (C) The ratio of Firefly to Renilla luciferase was used for quantification.

**Figure S2.**
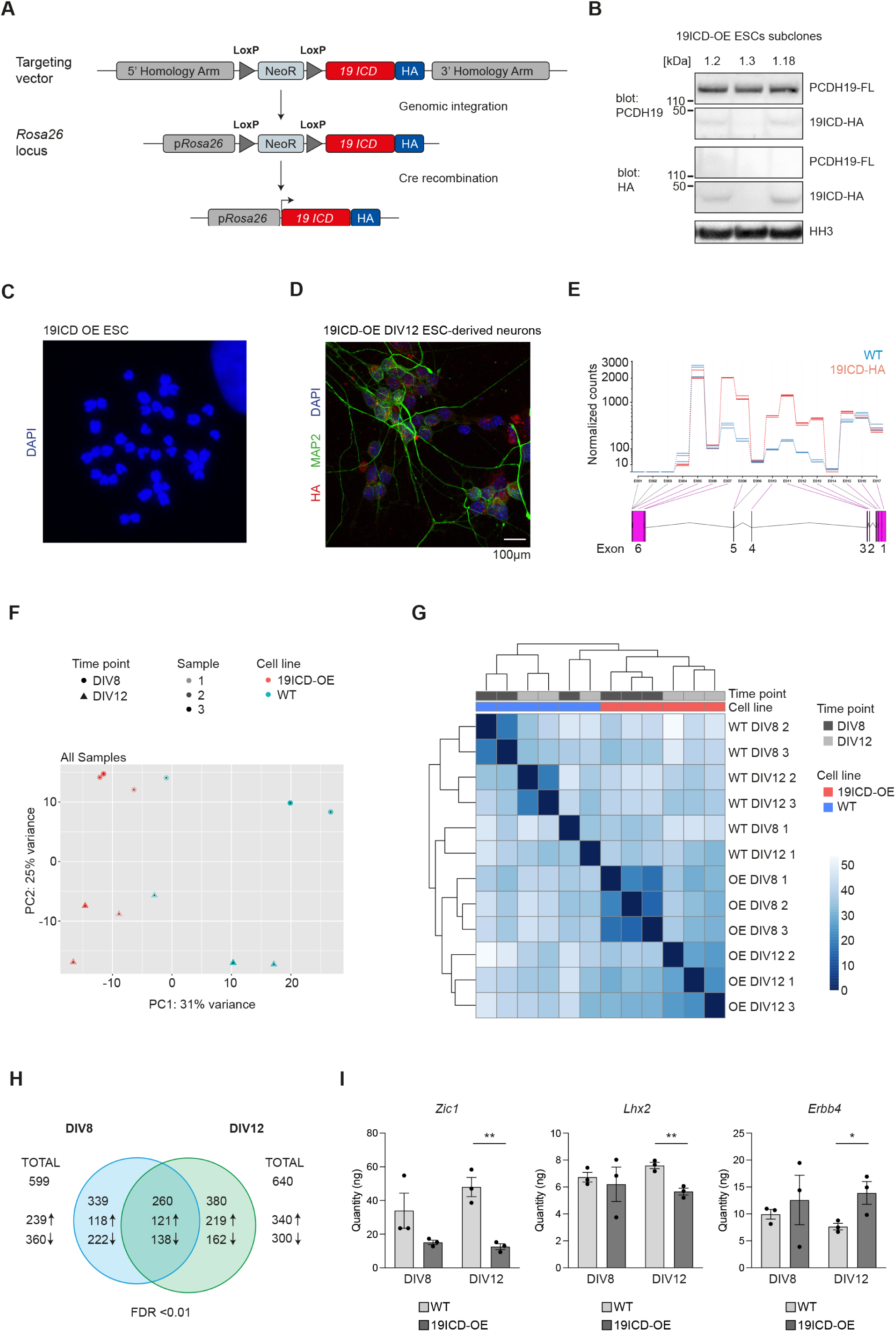
Generation of a new mESC-line to investigate the nuclear function of PCDH19-ICD. A. Schematic of the targeting of the *Rosa26* locus to express PCDH19-ICD-HA. B. WB of mESCs subclones, showing successful (1.2 and 1.18) and unsuccessful (1.3) targeting of the *Rosa26* locus. All clones express PCDH19, but only 1.2 and 1.18 show expression of PCDH19-ICD-HA. C. Representative karyotype of a targeted mESC subclone. Chromosomes were visualized with DAPI. D. DIV12 Neurons derived from 19ICD-OE mESCs stained for MAP2 (green) and HA (red). Nuclei are counterstained with DAPI (blue). Scale bar: 20 μm. E. *Pcdh19* exon usage. Normalised counts plotted along genomic coordinates of *Pcdh19* in DIV8 mESC-derived neurons (blue: WT (N=3), red: 19ICD-HA (N=3)). F. Principal component analysis on RNAseq results of DIV8 and DIV12 neuronal samples determined with DESeq2. Shapes indicate time points (DIV8, DIV12), colours indicate cell-lines (WT, 19ICD-OE) and alpha scale indicates samples belonging to each of the three independent differentiations (1, 2, 3). G. Correlation matrix showing hierarchical clustering of samples based on expression of all genes, determined with DESeq2. Blue scale represents distance between samples, with darker blue corresponding to smaller distances. H. Expression levels of differentially expressed genes in mESC-derived neurons at DIV8 and DIV12, determined by RT-PCR (N=3). Data information: cDNA quantity was obtained using a standard curve. For each of the 3 samples obtained from independent differentiations, 3 technical replicates were run, and their results were averaged. Data are presented as mean ± SEM. N: biological replicates (independent differentiations). Relevant P-values are indicated: *p < 0.05, **p < 0.01. Statistical analysis: two-tailed Mann-Whitney test (*Zic1* DIV8); unpaired two-tailed Student’s t-test (rest).

**Figure S3.**
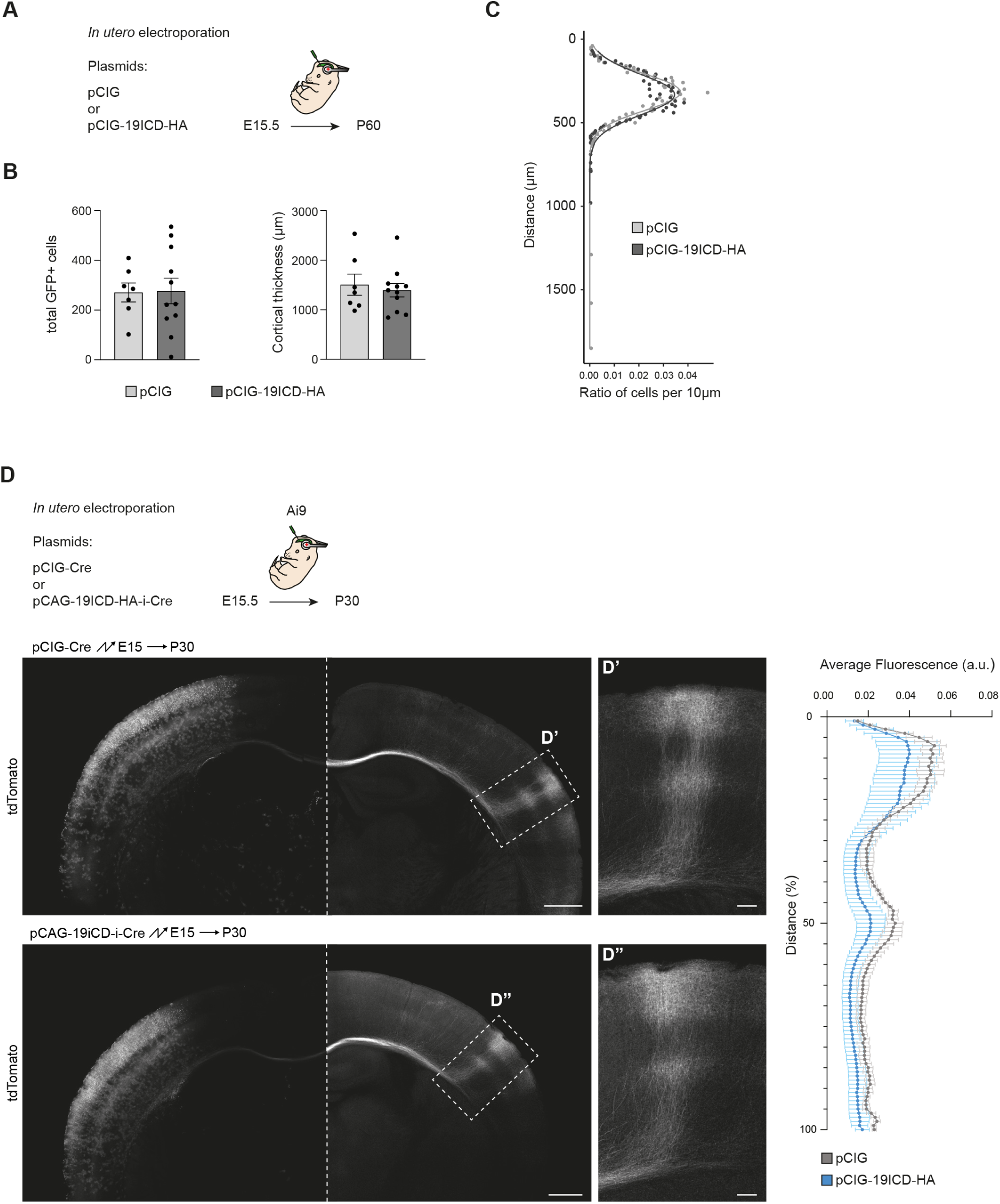
Neuronal number, position or axonal trajectories of upper layer cortical neurons are not affected by *in vivo* overexpression of PCDH19-ICD. A. Strategy for the *in utero* electroporation to express PCDH19-ICD in upper layer cortical neurons. B. Quantification (mean ± SEM) of the total number of electroporated neurons (top) and cortical thickness (bottom) for pCIG (N=7) and pCIG-19ICD-HA (N=11) brains. Average EP cells, pCIG: 271 ± 38, n = 7; pCIG-19ICD-HA: 277 ± 51, n = 11; unpaired t-test, *p* = 0.933; average cortical thickness, pCIG: 1505.41 ± 213.31 μm, n = 7; pCIG-19ICD-HA: 1394.79 ± 135.99 μm, n = 11; unpaired t-test, *p* = 0.6512. C. Kernel density analysis of the distribution of electroporated cells for pCIG and pCIG-19ICD-HA electroporated brains. 2-sample Kolmogorov-Smirnov test; D = 0.1622; P = 0.7154. D. *In utero* electroporation to analyse the effect of PCDH19-ICD overexpression on axonal development. Strategy (top) and representative confocal images of electroporated brains for the control (pCIG-Cre) and experimental (pCAG-19ICD-HA-ires-Cre) conditions. The right half of each image has been enhanced to allow visualization of the axonal arborizations. Higher magnifications of the contralateral hemisphere (D’, D’’) are also shown. Average fluorescence intensity (in arbitrary units) is shown for each of 100 bins spanning from the pial surface to the white matter for pCIG-Cre (N=3) and pCAG-19ICD-HA-ires-Cre (N=4). Scale bars: 500 μm for whole cortex, 100 μm for magnifications. The mixed effects model analysis applied to the data identified an effect of the distance to the pial surface on fluorescence intensity regardless of plasmid (mixed effects model; distance: F (1.219, 6.084) = 12.58; P = 0.0102, which reflects the two areas of axonal arborization in layers 2/3 and 5. However, there was no significant effect from the plasmid or the interaction between distance and plasmid (mixed effects model; plasmid: F (1, 5) = 0.7153, P = 0.4363; distance x plasmid: F (99, 494) = 0.3081, P > 0.9999. Data information: N: Biological replicate (electroporated brain). (B) Statistical analysis: unpaired two-tailed Student’s t-test. No statistically significant differences identified. (C) Kernel density analysis was performed on pooled data for N=7 and N=11 electroporated brains for pCIG and pCIG-19ICD-HA, respectively. Statistical analysis: Kolmogorov-Smirnoff test between histogram distributions of neuronal positions using 50 μm bins. No statistically significant differences identified. (D) Statistical analysis: mixed effects model with no assumption of sphericity.

